# Norovirus infection results in assembly of virus-specific G3BP1 granules and evasion of eIF2α signaling

**DOI:** 10.1101/490318

**Authors:** Michèle Brocard, Valentina Iadevaia, Philipp Klein, Belinda Hall, Glenys Lewis, Jia Lu, James Burke, Roy Parker, Alessia Ruggieri, Ian G. Goodfellow, Nicolas Locker

## Abstract

During viral infection, the accumulation of RNA replication intermediates or viral proteins imposes major stress on the host cell. In response, cellular stress pathways can rapidly impose defence mechanisms by shutting off the protein synthesis machinery, which viruses depend on, and triggering the accumulation of mRNAs into stress granules to limit the use of energy and nutrients. Because this threatens viral gene expression, viruses need to evade these pathways to propagate. Human norovirus is responsible for gastroenteritis outbreaks worldwide. Previously we showed that murine norovirus (MNV) regulates the activity of eukaryotic initiation factors (eIFs). Here we examined how MNV interacts with the eIF2a signaling axis controlling translation and stress granules accumulation. We show that while MNV infection represses host cell translation, it results in the assembly of virus-specific granules rather than stress granules. Further mechanistic analyses revealed that eIF2a signaling is uncoupled from translational stalling. Moreover the interaction of the RNA-binding protein G3BP1 with viral factors together with a redistribution of its cellular interacting partners could explain norovirus evasion of stress granules assembly. These results identify novel strategies by which norovirus ensure efficient replication propagation by manipulating the host stress response.

## INTRODUCTION

The synthesis of viral proteins, whose functions are essential to viral replication and propagation, is wholly dependent on gaining control of the host cell translation machinery. However, the accumulation of RNA replication intermediates or viral proteins imposes major stress on the host (1). In response to this stress, infected cells can induce several defence mechanisms to promote cell survival and limit the use of energy and nutrients, which can culminate in a global reduction of protein synthesis, while paradoxically allowing the rapid synthesis of antiviral proteins (1–3). Therefore, to replicate and spread, viruses need to balance their dependency on the host cell translation machinery with the adverse effect of antiviral proteins being synthesised by infected cells. As a consequence, interference with host mRNA translation represents a frequent evasion strategy evolved by viruses to subvert nearly every step of the host cell translation process (3,4). Most translational arrest strategies target translation initiation. This can be achieved by interfering with the initial cap-binding step mediated by eukaryotic initiation factor (eIF) 4F, targeting its integrity with viral proteases or its activity by modulating the mTOR or MAPK signaling pathways, or by altering the availability of the eIF2-GTP-tRNA_i_^Met^ ternary complex by phosphorylating eIF2α (3,4). eIF2α phosphorylation is mediated by four kinases (5–9). Among the four eIF2α kinases, protein kinase R (PKR) is activated by viral doublestranded RNA (dsRNA) sensing in the cytoplasm, however the other kinases: heme regulated eIF2α kinase (HRI), general control nonderepressible 2 (GCN2) and PKR-like ER localised kinase (PERK) can also be activated by infection cues such as oxidative stress, amino acid starvation and ER stress, respectively.

As consequence of translation initiation stalling and polysome disassembly, stalled mRNA initiation complexes can accumulate into membrane-less mRNA-protein granules called stress granules (SGs) (10–12). While it remains poorly understood this is a process driven by aggregation prone RNA binding proteins such as Ras-GTPase activating SH3 domain binding protein 1 (G3BP1) and T cell internal antigen-1 (TIA-1), driving protein-protein and RNA-protein interactions that result in a liquid-liquid phase transition (LLPT) and assembly of SGs (13). In addition, recently, RNA-RNA interactions have also been demonstrated to contribute to SG assembly (14,15). SGs are highly dynamic, able to rapidly assemble to act as storage sites for mRNAs, fuse and then dissolve upon stress resolution. The protein composition of SGs is highly variable and alters depending upon the type of stress induction (16–18). In addition, while the bulk content of cytoplasmic mRNAs can be sequestered within SGs, specific exclusion of transcripts allows for stress-specific translational programme to combat stress (19,20). Furthermore SG or specific antiviral SGs (avSG) have been proposed to play a role in antiviral signaling as key signaling proteins including MDA5 and PKR are known to localise to SGs and SG formation is involved in PKR activation (21,22). Although some may exploit SG for their replication many viruses have evolved strategies to antagonize SGs, by impairing the eIF2α sensing pathway or through the cleavage and sequestration of SG-nucleating proteins (23,24). Among these, G3BP1 is prime target for several viruses as it is proteolytically cleaved by enterovirus and calicivirus proteases, sequestered by the alphavirus nsp3 protein or repurposed by the Dengue virus RNA and during vaccinia infection (23,24). Therefore viral infections have a profound impact on translational control, acting at the nexus between translation stalling and SG assembly.

The *Caliciviridae* family comprises small non-enveloped positive-strand RNA viruses of both medical and veterinary importance. Among these, human norovirus (HuNoV) is a leading cause of acute gastroenteritis and food-borne illnesses worldwide, accounting for nearly one fifth of cases and 200,000 deaths per annum (25,26). This also results in a significant economic burden to the health organisations with an estimated economic impact of $60 billion (27,28). The genogroup GII genotype 4(GII.4) strains are responsible for the majority of outbreaks, including pandemics. While the diarrhoea and vomiting symptoms are acute and self-resolving in most children and adults, HuNoV has been reported to cause persistent infections, sometimes fatal, in the very young and elderly populations (29–31) and HuNoV infections exacerbates inflammatory bowel disease or has been associated with neonatal enterocolitis (32,33). Currently no specific vaccine or antiviral has been approved, and efforts to develop these have been hampered by the difficulty in culturing HuNoV *in vitro* and the lack of robust small animal models (34,35). Recent studies have demonstrated that immortalised B cells and stem-cell derived intestinal organoids support HuNoV replication in culture, however these experimental systems are technically challenging and currently lack the degree of robustness required for a detailed analysis of the viral life cycle (35–37). However, the related calicivirus murine norovirus (MNV) can be propagated in cell culture, has reverse genetics systems and small animal model and remains the main model for understanding the life cycle of caliciviruses (38,39). MNV and HuNoV share many characteristics such as their genome structure, environmental stability, faecal–oral transmission, replication in the gastrointestinal tract, and prolonged viral shedding (39). Recently, the cell surface protein CD300lf was identified as a proteinaceous receptor for MNV (40). Moreover, the expression of CD300lf alone in human cells is sufficient to enable MNV replication, suggesting the intracellular replication machinery for norovirus is conserved across species. MNV possesses a genome of ~7.5kb in length, capped by a viral genome-linked protein (VPg) covalently attached at the 5’ end and polyadenylated at the 3’ end. The VPg protein directs the translation of the MNV ORF1-4 by interacting directly with eIFs (41,42) generating polyproteins subsequently proteolytically processed into three structural and seven non-structural proteins (43). Meanwhile our previous studies have shown that MNV infection regulates translation in several ways by controlling the activity of multiple eIFs by inducing eIF4E phosphorylation via the MAPK pathway, and by cleavage of PABP and eIF4G by the viral protease or cellular caspases (41,44). Therefore, MNV induces hosts responses that affect translation and could coordinate SG assembly. Furthermore, while infection with another norovirus model, Feline Calicivirus (FCV) disrupts the assembly of SGs by inducing G3BP1 cleavage, MNV infection has no impact on G3BP1 integrity (45). Herein, we address the complex interaction between MNV, the control of the host cell translation machinery and the SG response pathway. We show that while MNV infection impairs host cell translation early post-infection, this translational suppression is uncoupled from the activation of the eIF2α-dependent stress response and SG assembly. Instead viral proteins interact with G3BP1 resulting in the assembly of G3BP1-containing granules that differ markedly from arsenite-induced granules. This provides evidence that MNV has evolved an evasion strategy to counteract the cellular SG response during infection.

## MATERIAL AND METHODS

### Cells, retroviral transduction and MNV-1 production

Murine macrophage cells RAW264.7 and microglial cells BV2 were maintained in Dulbecco’s modified Eagle’s medium (DMEM), 4,5g/L D-glucose, Na Pyruvate, L-Glutamine supplemented with 10% foetal calf serum (FCS), 100 U of penicillin/mL, 100μg of streptomycin/mL, 10mM HEPES, and 2mM L-glutamine (all supplements purchased from Invitrogen) at 37°C in a 5% CO2 environment. Primary murine bone marrow derived macrophages (BMDMs) were prepared as described in (46) from C57BL/6 mice bones and cultivated in the same medium supplemented with 50ng/ml of M-CSF (Preprotech). BV2 GFP-G3BP1 Puro, BV2 mCherry-eIF3E Neo, BV2 GFP-G3BP1/mCherry-eIF3E Puro/Neo cells and control cells BV2 Neo and BV2 Neo/Puro cells were produced by retroviral transduction of BV2 cells as described in (47). Transduced cells were selected in complete medium at 2μg/ml of Puromycin (Sigma) or 100μg/ml G418 or 2μg/mL of puromycin and 100μg/ml G418 (Sigma) and propagated in complete medium containing 1μg/mL of puromycin or, 50μg/mL G418 or 1μg/mL of puromycin and 50μg/mL G418. For live-cell imaging, high expressing cells were sorted by flow cytometry. MEFs w.t. and S51A – CD300lf were produced by retroviral transduction of MEFs w.t. and S51A (kind gift from M. Coldwell, University of Southampton) with the lentiviral particles containing the full-length CD300lf sequence and hygromycin resistance gene, selected at 100μg/ml of hygromycinB (Sigma) and propagated in complete medium containing 50μg/ml of hygromycinB. Murine norovirus 1 (MNV-1) strain CW1 was described previously (38). MNV-1 was propagated in BV2 cells as described in (45) with an extra step of concentration using Amicon Centrifugal Filters 100k (Millipore). Virus titres were estimated by determination of the TCID50 in units per millilitre in RAW264.7 cells and infections were carried at a MOI of 10 unless stated otherwise. The times post-infection refer to the time elapsed following medium replacement after a 1h inoculation period.

### SG formation, p-eIF2α induction and drug treatment

SG were induced using arsenite (Sigma) at a final concentration of 0.1mM for 1h at 37°C except for MEFs (0.5mM for 1h) and the disassembly was forced by adding cycloheximide (Sigma) at 10μg/ml 30min after the arsenite treatment. Phosphorylation of eIF2α was induced either by treatment with arsenite as described above, tunicamycin (Sigma) at 5μg/ml for 6h or by UV irradiation at 20mJ/cm^2^ using a Stratalinker 1800 (Stratagene). ISRIB (Sigma) was added to the cells at a final concentration ranging from 10 to 1000nM and A-92 (Axon Medchem) from 100 and 1000nM at t=0hp.i and for the indicated times.

### Immunoblotting

Immunoblotting analysis was performed as described in (45). Briefly, approximately 10^^6^ BV2 cells or BMDMs, 2×10^^6^ RAW264.7 cells or 5.10’^5^ MEF cells were plated onto 35mm dishes and infected the next day with MNV at a MOI of 10 as described above. At the indicated times, cells were lysed in 150μL of 1x Gel Loading Buffer (New England Biolabs), sonicated and boiled 5min at 95°C. Cell lysates were separated by SDS-PAGE, using 10μg of total proteins, and transferring the proteins to nitrocellulose or polyvinylidene difluoride membranes. These were then probed with the following primary antibodies: rabbit anti-eIF2α (1:1,000, Cell Signaling), mouse anti-P-eIF2α (1:5,000, Invitrogen), goat anti-eIF3B (1:2,000, Santa Cruz), rabbit anti-NS3 (1:1,000, kind gift from I. Goodfellow), rabbit anti-NS7 (1;10,000), rabbit anti-G3BP1 (1:2,000, Sigma), rabbit anti-Caprin1 (1:1,000, Bethyl Laboratories), rabbit anti-USP10 (1:1,000, Bethyl Laboratories), rabbit anti-rpS6 (1:1,000, Cell Signaling), mouse anti-GAPDH (1:20,000, Invitrogen); followed by incubation with the appropriate peroxidase labelled secondary antibodies (Dako) and chemiluminescence development using the Clarity Western ECL Substrate (Bio-Rad). The results were acquired using the VILBER imaging system. Phos-tag analysis was performed to detect mobility shift of phosphorylated eIF2α as described in (48). Briefly, 2×10^^6^ RAW264.7 cells were scraped on ice at different times post infection, pelleted for 2min at 4°C full speed, washed in PBS and lysed in 125μl of cold Lysis Buffer (Tris HCl pH 7.4 50mM, NaCl 150mM, Triton X100 1%) supplemented with antiphosphatase and antiprotease cocktails (Roche). Lysates were incubated for 30min on ice before being centrifuged at 13K rpm for 30min at 4C and the supernatants transferred to new Eppendorf tubes. Protein concentrations were measured by BCA (Pierce) according to the manufacturer’s instructions. Twenty-five μg of total proteins were denatured 3min at 95°C and loaded onto Mn^2+^-Phos-tag SDS-PAGE gel prepared as described in (49,50). Both basal and phosphorylated forms of eIF2α were visualised using an antibody anti-eIF2α. Signal was detected using the Advance ECL Chemocam Imager (Intas Science Imaging) and band quantified using LabImage 1D Software (Intas Science Imaging).

### Immunofluorescence microscopy

10^^6^ BV2 cells and BMDMs or 2×10^^6^ RAW264.7 cells were plated on glass coverslips in a 6 wells plate, 5×10^^5^ MEF w.t. and S51A were plated on poly-L-Lysine coated glass coverslips according to the manufacturer instructions (Sigma). Cells were infected the next day with MNV at MOI of 10 for the indicated times, fixed with a solution of 4% PFA (Sigma) in PBS for 30min at room temperature (RT), washed in PBS and store at 4°C. After permeabilisation with 0.1% Triton-X100 (Sigma) in PBS for 5min at RT and 3 washes with PBS, the coverslips were blocked with 0.5% BSA (Fisher) in PBS for 1h at RT then incubated with 200μl of primary antibodies solution for 2h at RT. After 3 washes with PBS, coverslips were incubated in the dark with 200μl of fluorescently-labelled secondary antibodies solution containing 0,1μg/mL DAPI solution (Sigma). The coverslips were then washed and mounted on slides with a drop of Mowiol 488 (Sigma). Confocal microscopy was performed on a Ti-Eclipse - A1MP Multiphoton Confocal Microscope (Nikon) using the Nikon acquisition software NIS-Elements AR. Primary antibodies dilutions: rabbit anti-NS3 (1:600), mouse anti-dsRNA (1:1,000), mouse anti-puromycin (1:5, http://dshb.biology.uiowa.edu/PMY-2A4), mouse anti-G3BP1 (1:400, Invitrogen), goat anti-eIF3B (1:400, Santa Cruz). Secondary antibodies were all purchased from Invitrogen: goat antirabbit Alexa 488, goat anti-mouse Alexa 555, chicken anti-mouse Alexa 488, donkey anti-goat Alexa 555 and chicken anti-rabbit Alexa 647. All quantification analysis were made using Image J software package Fiji (http://fiji.sc/wiki/index.php/Fiji) and all images were processed using the Nikon analysis software NIS-Element AR.

### Ribopuromycylation assay and quantification of fluorescence intensities

Quantification of *de novo* protein synthesis was performed as described by (51). Briefly, cells were plated on coverslip in 6 wells plate and infected the next day with MNV at MOI of 10 for the indicated times. Prior fixation, cells were treated with 10μg/ml of puromycin (Sigma) for 2 min at RT to label the nascent polypeptide chains before addition of 180μM of emetin (Sigma) to block the translation elongation with a further incubation of 2 min at 37°C. Coverslips were washed 3 times in prewarmed complete medium, fixed with 1ml of 4% PFA in PBS for 15min at RT, washed 3 times in PBS. Cells treated with 100μg/mL cycloheximide for 5 min prior to puromycylation were used as control. Cells treated with 5μg/mL tunicamycin for 6h prior to puromycylation were used as a control of P-eIF2α-dependent inhibition of translation. Fluorescence intensities were quantified by using Image J software package Fiji as described in (48).

### Live-cell imaging

MNV-1 was produced has described above except BV2 cells were cultivated in microscopy medium (phenol red free DMEM, 4,5g/L glucose supplemented with 10% FCS, 100U/ml of penicillin, 100μg/mL of streptomycin, 10mM HEPES, and 2mM L-glutamine and 1x Na Pyruvate (Invitrogen)).

10^^5^ BV2 GFP-G3BP1/mCherry-eIF3E Neo Puro cells were seeded on 12-well glass-bottom plates (Cellvis) in 2ml microscopy medium (phenol red free DMEM, 4,5g/L glucose, HEPES, L-glutamine (Invitrogen) supplemented with 10% FCS, 100U/ml of penicillin, 100μg/mL of streptomycin). Cells were infected with MNV-1 at a MOI of 20 and directly transferred for live-cell imaging. Images were acquired with a 40x magnification (Nikon objective CFI P-Fluor 40x N.A. 1.30 oil immersion) every 15 min for 20 h using an UltraVIEW VoX Spinning Disk Confocal Microscope from PerkinElmer (Volocity software package) and a Hamamatsu Orca Flash 4 camera.

### Biotin-isoxazole (b-isox) induction liquid liquid phase transition

Treatment were performed as described in (52). Briefly, 2×10^^7^ RAW264.7 cells were plated onto 10cm dishes, Mock or MNV-infected the next day at a MOI of 10 and further cultivated for 10h post infection (p.i). The dishes were placed on ice and washed with cold PBS (from 10x solution – Lonza) and the cells lysed on ice with 1ml of EE buffer (Hepes pH 7.4 50mM, NaCl 200mM, Igepal 0.1%, EDTA pH8 1mM, EGTA pH8 2.5mM, Glycerol 10%, DTT 1μM supplemented with RNAsin (Promega) and anti-protease cocktail (Roche)). Lysates were transferred to a 1.5ml Eppendorf and incubated with agitation for 20min at 4°C before being spun down at 13,000 rpm 15min at 4°C. 50μl of the supernatant was kept as “input”, mixed with an equal volume of 2x Loading buffer (Cell Signaling) and boiled at 95°C for 5min. The remaining supernatant was supplemented with 100μM of b-isox (Sigma) and the precipitation reactions were carried out for 90min at 4°C with agitation. Aggregates were pelleted down by centrifugation at 10,000xg for 10min at 4°C. 50μL of the supernatant was kept as “soluble fraction”, mixed with an equal volume of 2x loading buffer and boiled at 95°C for 5min. The pellets were washed twice with 500μL of cold EE buffer, spin down at 10,000g 10min at 4°C, resuspended into 200μL of 1x Gel Loading Buffer (New England Biolabs) as “insoluble fraction” and boiled at 95°C for 5min.

### Immunoprecipitation

10^^7^ BV2 GFP-G3BP1 cells were plated onto 10cm dishes, infected the next day with MNV or MNV(UV) at a MOI of 10 and further cultivated for 6 and 9hp.i. The dishes were placed on ice and washed twice with cold PBS (from 10x solution – Lonza) and the cells lysed on ice with 500μL of cold Lysis Buffer (Tris-HCl pH7.4 50mM, NaCl 50mM, MgCl_2_ 15mM, CHAPS 0.12% (w/v), RNAsIn (Promega) 200u/ml, DTT 2mM, Beta-Glycerophosphate 1,75mM, NaF 50mM, NaPyrophosphate 2mM, antiprotease cocktail (Roche). Lysates were incubated 30min on ice, centrifuged at 2krpm, 2min at 4°C, the supernatant transferred to a new Eppendorf tube and flash frozen in liquid nitrogen. 25μL of magnetic Sepharose-Protein G beads (Invitrogen) per sample were washed twice in cold PBS and incubated overnight in IP buffer (Tris-HCl pH7.4 50mM, NaCl 100mM, MgCl_2_ 15mM, yeast tRNA 100μg/mL, BSA 5ug/mL, RNAsin 200u/mL (Promega) DTT 2mM, antiprotease cocktail (Roche) at 4°C with either 0.5μg of anti-GFP antibody (Invitrogen, Monoclonal mAb 3E6) or 0.5μg of normal mouse IgG (Santa Cruz). The beads were washed 3 times in IP buffer, resuspended in 250μL of IP buffer before adding 250μg of lysate at 1mg/ml to each tube. The IgG-beads:lysates mixes were incubated for 1hr at 4°C on a rotary wheel, washed 3 times in IP buffer before being resuspended into 100μL of 1x Gel Loading Buffer (New England Biolabs), boiled at 95C for 5min and the supernatants transferred to new Eppendorf.

### G3BP1 interactome

9×10^^Ø^ BV2 GFP-G3BP1 Puro cells were infected with MNV (MOI 10 for 9h) or stressed with arsenite (0.1mM for 1h) then were crosslinked with 1% formaldehyde in PBS for 10 min at RT, and quenched by adding 125mM of glycine for 10 minutes at RT. The cells were spun at 4°C for 5 minutes at 230xg and the pellet was re-suspended in 1ml of SG lysis buffer (50mM Tris-HCl pH 7.4, 100mM Potassium Acetate, 2mM Magnesium Acetate, 0.5mM DTT, 50μg/mL Heparin, 0.5% NP-40, 1 complete mini EDTA-free protease inhibitor tablet (Roche)) and then lysed by passing through a 25G 5/8 needle seven times on ice and spun at 1,000xg for 5 minutes. The supernatant was collected and spun down at 18,000xg for 20 min at 4°C. Subsequently the supernatant was removed and the pellet was resuspended in 1mL of SG lysis buffer and again spun down at 18,000xg for 20 min at 4C. Finally the pellet was re-suspended in 300μL of SG lysis buffer (stress granule core enriched fraction) and nutated at 4°C for 1hr with 60μl of magnetic Dynabeads Protein A (Invitrogen). Once the beads have been removed, the supernatant was incubated with specific antibody (1μg of rabbit anti-GFP antibody and 1μg of IgG) and nutated overnight at 4°C. The unbounded antibody was removed by centrifugation at 18,000xg for 20 min at 4°C, and the pellet was re-suspended in 300μL of SG lysis buffer before to incubate it with 60μL of Dynabeads Protein A at 4°C for 3h. The Dynabeads were then washed for 2 min at 4°C with 1mL of wash buffer 1 (SG lysis buffer and 2M Urea), for 5min at 4°C with 1mL wash buffer 2 (SG lysis buffer and 300mM potassium acetate), for 5min at 4°C with 1mL SG lysis buffer and seven times with 1ml of TE buffer (10mM Tris HCl pH 8.0, 1mM EDTA pH 8.0). Mass spectrometric analysis by LC-MS Orbitrap of the Dynabeads was performed at the Proteomics Facility at University of Bristol as detailed in the Supplementary Data.

### Statistical Analyses

Statistical analysis were performed by using the GraphPad Prism software. Statistical significances were calculated by performing a two-tailed Student’s t test (****, p<0.0001, ***, p<0.001, **, p<0.01, n.s., not significant).

## RESULTS

### MNV infection results in hallmarks of translational stalling

Previous studies have demonstrated that the activity of translation initiation factors, such as eIF4E or PABP, is regulated during MNV infection, resulting in global reduction in translation (44,53). To better understand the dynamic control of translation within infected cells, we addressed the global translational efficiency using single cell analysis by measuring the incorporation of puromycin, a tRNA structural mimic which specifically labels actively translating nascent polypeptides and causes their release from ribosomes (51,54). Anti-puromycin antibodies can then be used to detect puromycylated native peptide chains by confocal microscopy. To account for different cellular models used to study MNV replication, we chose to address this event in the murine macrophages cell lines RAW264.7 and primary murine bone marrow derived macrophages (BMDMs). Quantification of the puromycin signal intensity showed a significant decrease of protein synthesis following treatment with the translation elongation inhibitor cycloheximide (CHX) (55) in both RAW264.7 and BMDM cells (Figure 1 A and B and Figure 1 D and E). RAW264.7 and BMDM cells were infected with MNV and fixed at 6, 9 and 12h p.i. or 9 and 15h p.i, respectively. Quantification of the puromycin signal in individual infected cells, defined by immunostaining of MNV non-structural protein NS3, showed a progressive reduction in protein synthesis over time. In RAW264.7 and BMDM cells this decrease was detectable from 9h p.i onward. This time point correlated with peak viral protein production in direct comparison to cells infected with UV-inactivated MNV (MNV(UVi)) as a non-replicative control as evidenced by immunoblotting of RAW264.7 (from 2 to 10h p.i.) and BMDM (from 3 to 15h p.i.) cells (Figure 1 C and F). These results confirm that infection leads to a translation shut-off in cell lines and primary cells permissive to MNV replication and further extends our previous observations (44).

**Figure 1.**
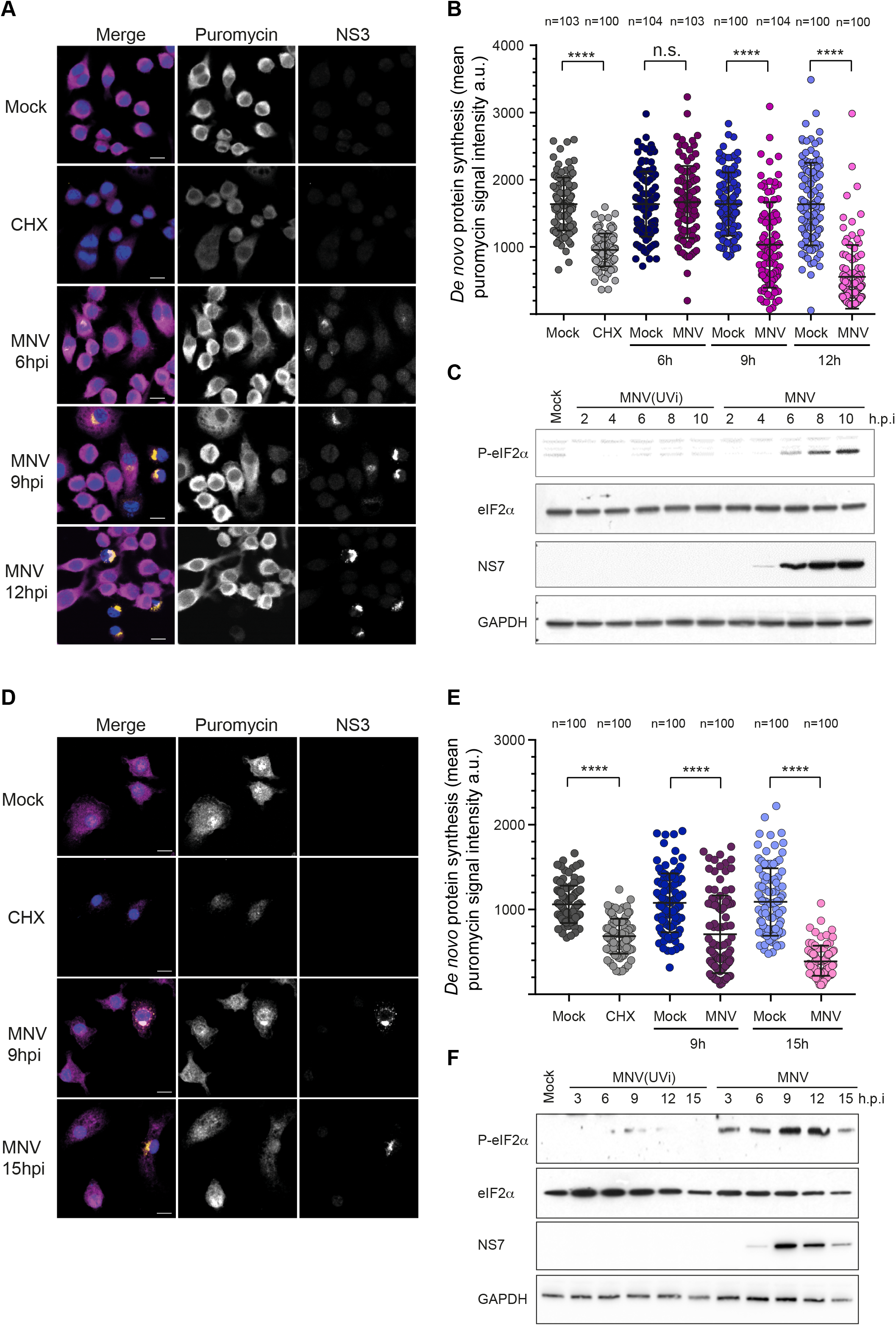
MNV replication results in gradual translational shut-off and hallmarks of eIF2α signaling activation in cell lines and primary cells. Decrease protein synthesis in MNV-infected RAW264.7 cells (A and B) or primary murine BMDMs (D and E). Naïve cells (Mock) or MNV-infected cells (MOI 10) for the indicated times were incubated with 10μg/ml of puromycin to label the nascent polypeptidic chains prior to fixation. CHX-treated cells were used as a negative control. Puromycin-labelled chains were visualised by immunostaining against puromycin (magenta) and infected cells were detected by immunostaining against MNV NS3 (gold). Nuclei were stained with DAPI. Representative views are shown for RAW264.7 cells (A) and BMDMs (D). Scale bars, 10μm. Representative scatter plots of *de novo* protein synthesis measured by fluorescence intensity of the puromycin signal (n=3) in RAW264.7 cells (B) and BMDMs (E), a.u. arbitrary units. Statistical significance and number of analysed cells (n) are given at the top. **** *P* < 0.0001. MNV infection induces an increase of the phosphorylation levels of eIF2α (C and F). Representative Western Blot analysis (n=3) of RAW264.7 cells (C) and BMDMs (F) infected (MOI 10) for the indicated times with replicative MNV or non-replicative virus MNV(UVi). Naive cells (Mock) were cultivated in parallel for 10h (C) or 15h (F).

During viral infection, the accumulation of double-stranded (ds) RNA replication intermediates commonly leads to the activation of the cytoplasmic pattern recognition receptor PKR, which results in the downregulation of the host translation machinery via phosphorylation of the translation initiation factor eIF2α, a hallmark of viral sensing (9). Consequently, we then addressed the phosphorylation status of eIF2α in MNV-infected cells. Time course analysis by immunoblotting showed an accumulation of P-eIF2α in MNV-infected cultures concomitant with the accumulation of viral products and translational shut-off but not in the cells inoculated with the UV-inactivated virus (Figure1 C and F). This result suggests that MNV replication triggers the expected host response to viral infection, through the activation of eIF2α phosphorylation, culminating in the host translation shut-off.

### MNV infection does not induce canonical stress granule assembly

The inhibition of translation initiation via eIF2α-dependent or independent pathways during infection is intimately linked to the assembly of SGs (1). To counteract this stress response, many viruses have evolved strategies to impair SG formation (24). We previously reported that FCV prevents SG formation through the cleavage of the nucleating factor G3BP1 by the viral proteinase NS6^Pro^, while we observed no impact of MNV infection on G3BP1 integrity in the J774A.1 murine macrophages cell line (45). To better examine a potential role for the activation of the stress response in MNV-induced translational suppression and to better understand the interaction of noroviruses with SGs, we analysed SG formation in more detail. (Figure 2).

**Figure 2.**
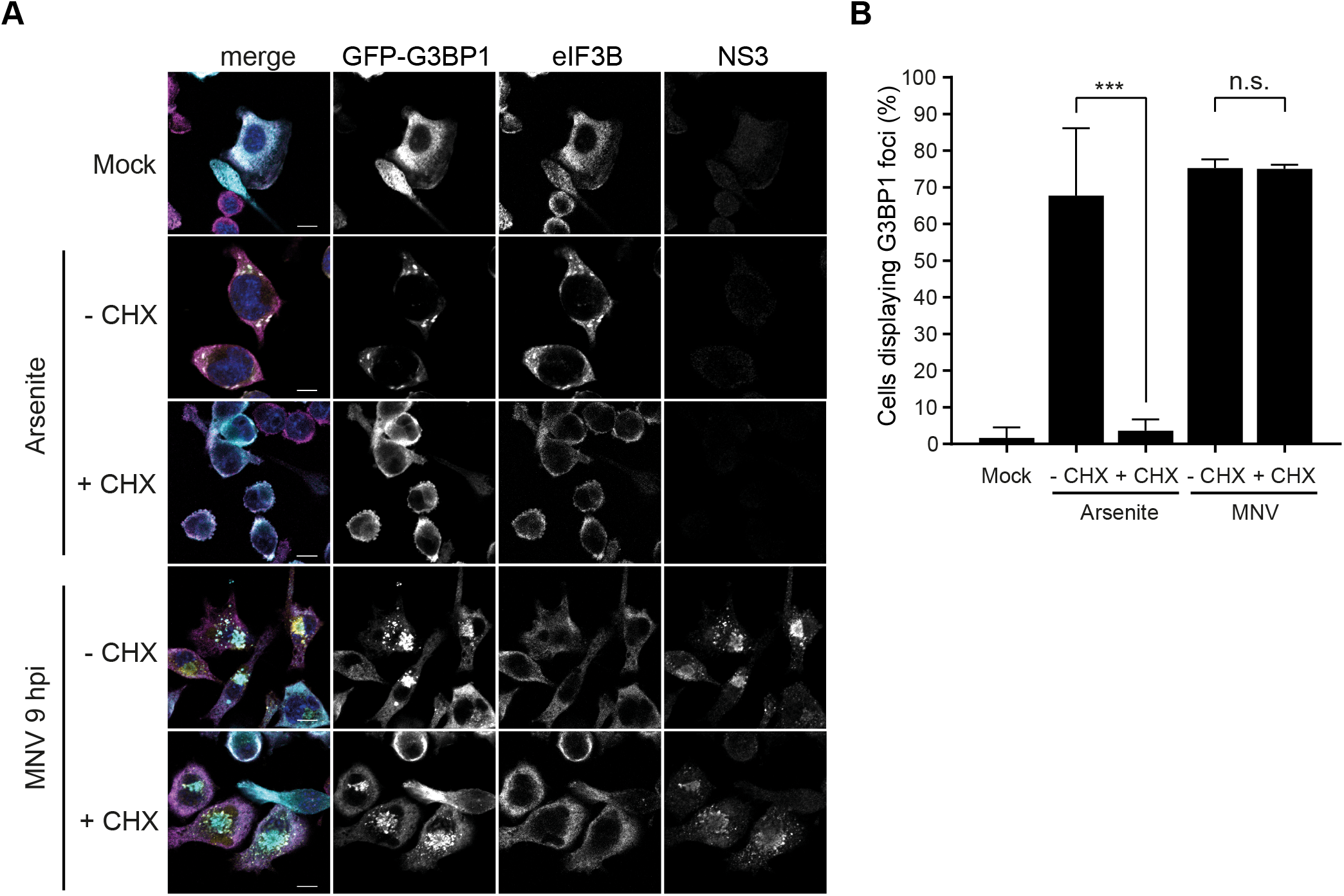
MNV infection results in the assembly of non-canonical cytoplasmic G3BP1 granules. BV2 GFP-G3BP1 cells were infected with MNV (MOI 10) for 9h p.i. arsenite-treated cells were used as positive control for SG formation. For forced SG disassembly, cells were treated with 10μg/ml of CHX for 30min (+CHX). (A) Confocal analysis of the formation of MNV-induced granules in GFP-G3BP1 (cyan) positive cells. Samples were stained for the SG marker eIF3B (magenta) and infected cells detected by immunostaining against MNV NS3 (gold). Scale bars, 10μm. (B) Representative bar plot (n=3) of the percentage of cells displaying GFP-G3BP1 foci, mean ± SD for 100 GFP-positive cells (Mock and arsenite) and GFP- and NS3-positive cells (MNV) analysed across at least 10 acquisitions. Statistical significance shown above the bars, ***, *P* < 0.001, n.s., not significant.

Most known SG markers, such as TIA-1, G3BP1 or Caprin1, were found unsuitable for immunofluorescence analysis in RAW264.7 cells (data not shown). Therefore for our studies we used the murine microglial cell line BV2 stably over-expressing an exogenous GFP-tagged G3BP1 (BV2 GFP-G3BP1), allowing us to detect the subcellular distribution of total G3BP1 by immunofluorescence microscopy. In addition to G3BP1, the eukaryotic initiation factor eIF3B was also used as a *bona fide* SG marker, and the viral protein NS3 as a marker for infected cells. First, we confirmed that this cell line was as permissive to MNV infection as the non-transfected parental line by assessing the viral titers following infection at a MOI of 1 (Figure S1A). While no SG were detected in uninfected cells, as expected, upon treatment of BV2 GFP-G3BP1 for 1h with 0.5mM arsenite, which induces oxidative stress and eIF2α-dependent inhibition of translation, both G3BP1 and eIF3B relocated to SGs (Figure 2A). In stark contrast, infection with MNV for 9h resulted in the accumulation of G3BP1 in large cytoplasmic foci localized closely to the nucleus, colocalising with an accumulation of the viral protein NS3. The foci identified were devoid of eIF3B suggesting that they did not represent canonical SGs. Noteworthy, these structures highly resemble previously characterized MNV replication complexes (56–58). To ensure that this was not an experimentally induced artefact as a result of GFP-G3BP1 over-expression, we confirmed that the endogenous G3BP1 redistributed in a similar manner (Figure S1B). Furthermore, another SG component eIF3E was excluded from MNV-induced G3BP1 foci in BV2 cells constitutively expressing both GFP-G3BP1 and mCherry-eIF3E (Figure S2).

SGs are assembled in response to the dynamic exchange of mRNAs transcripts between polysomes and SG. Therefore, the translation elongation inhibitor CHX, by trapping the ribosomes on mRNA transcripts, prevents the accumulation of stalled initiating complexes required for the formation of SGs and is used to discriminate between canonical SGs and other cytoplasmic granules (59). Treatment of BV2 GFP-G3BP1 cells with CHX resulted in a potent block of SG assembly upon treatment with arsenite (Figure 2A and 2B). In contrast, CHX treatment had no significant impact on the accumulation of G3BP1 foci at 9h p.i. and its co-localisation with NS3 (Figure 2A and 2B). Altogether, these results suggest that MNV infection leads to a redistribution of G3BP1 which is not driven by the accumulation of stalled initiating complexes, and may represent an assembly of virus-specific granules.

### G3BP1 ability to undergo SG-like phase transition is unaffected by MNV

Previous studies have demonstrated that G3BP1 is targeted by multiple viruses to control the formation and function of SGs (60,61). While the calicivirus proteinase NS6^Pro^ mediates G3BP1 cleavage during FCV infection, it does not impact G3BP1 integrity during MNV infection (45). The ability of G3BP1 to undergo LLPT, and drive SG assembly in cells under stress is dependent upon dynamic post-translational modifications. This has been linked in particular to the removal of methyl groups on Arginine residues (62) and the phosphorylation status of the Serine 149 (63). These modifications prevent unnecessary and untimely formation of G3BP1 aggregates by neutralizing the physical ability of G3BP1 to homo-(64) and heteropolymerise with other SG-associated proteins and non-translating RNAs. The ability of G3BP1 to undergo LLPT can by recapitulated *in vitro* by inducing selective condensation of the low-complexity aggregation-prone proteins with biotinylated-isoxazole (b-isox), in conjunction with an EDTA-EGTA lysis buffer releasing transcripts from polysomes (52,65). The b-isox-induced aggregates can then be isolated from the soluble fraction by centrifugation, respectively defined as pellet and supernatant.

We therefore addressed the possibility of that inhibitory post-translational modifications of G3BP1 in MNV-infected cells prevent LLPT by adding b-isox to cytoplasmic extracts of RAW 264.7 cells, mock-or infected with MNV for 10h (Figure 3). Quantification analysis of immunoblotting analysis showed no difference in the levels of G3BP1 recovered in the b-isox precipitated fractions (pellets) compared to the inputs in the extracts from mock-or MNV-infected cells. Moreover, the canonical SG marker Caprin1 and to a lesser extend the 48S component eIF3B and ribosomal protein rpS6 were also precipitated with G3BP1, suggesting both the precipitation of low-complexity proteins and the interaction of G3BP1 with non-translating mRNPs, which seems to confirm an authentic recapitulation of P-eIF2α-dependent SG assembly in this experimental set-up. Conversely, we did not observe precipitation of the viral RNA polymerase NS7. Overall, this suggests that MNV infection does not result in conditions that hinder G3BP1 ability to undergo the LLPT normally associated with the accumulation of SGs. Thus the condensation of G3BP1 in MNV-induced granules does not seem responsible for preventing SG formation by sequestering G3BP1.

**Figure 3.**
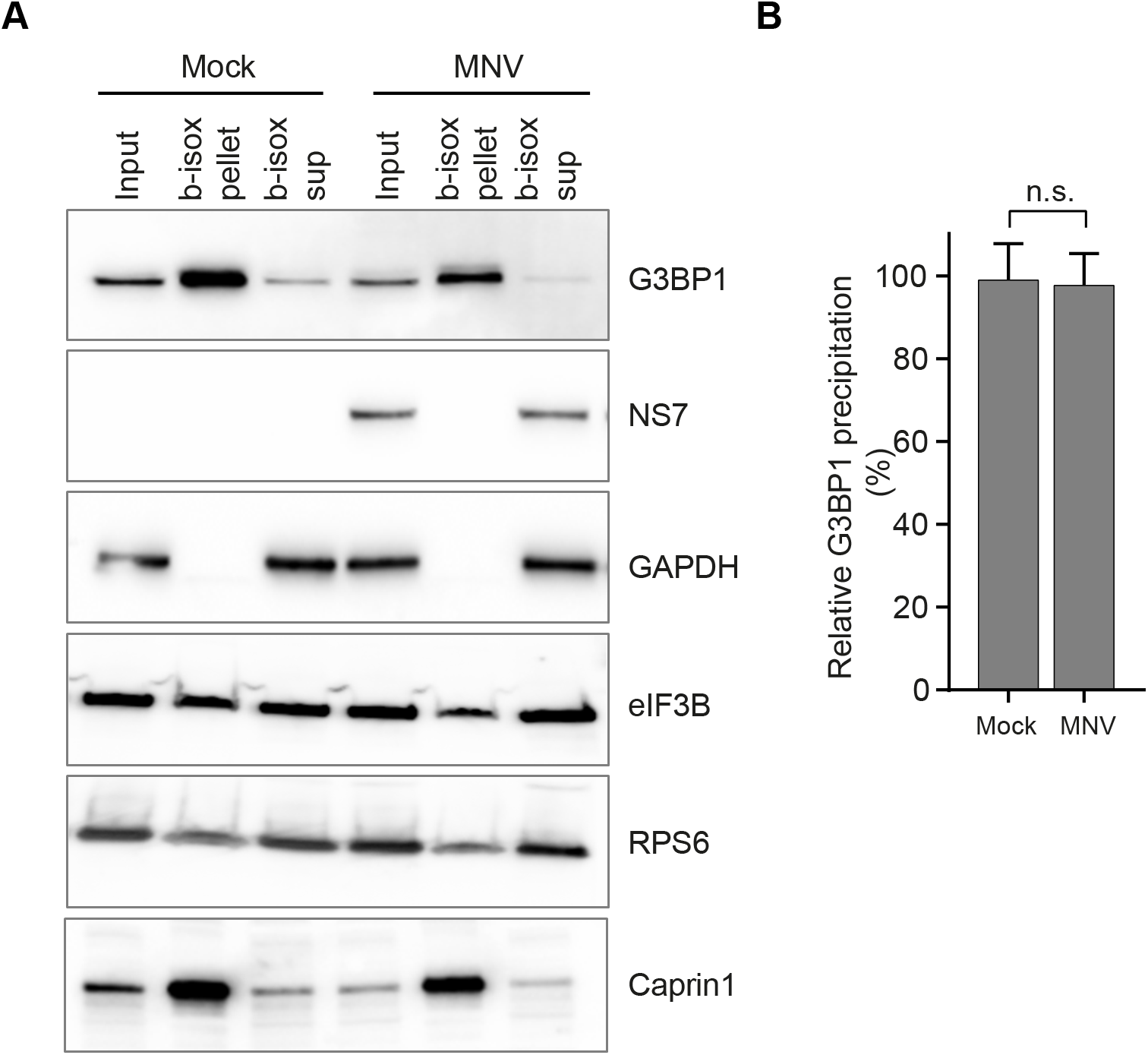
MNV infection does not prevent a LLPT-induced G3BP1 condensation. *In vitro* SG assembly induced by b-isox in RAW264.7 cells extracts (A and B). Naïve (Mock) or MNV-infected RAW264.7 cells for 10hp.i. (MOI 10) were lysed and 100μM of b-isox added to the extracts before centrifugal separation between b-isox induced SG-like aggregates (b-isox pellet) and supernatant (b-isox sup). (A) Western blot analysis of the distribution of cellular and viral proteins. For each condition, samples were loaded as Input, b-isox pellet and b-isox supernatant from left to right. (B) Bar plot of the relative G3BP1 precipitation as the ratio between the level of G3BP1 in the b-isox pellet compared to the Input for the Mock (-MNV) and MNV-infected (+MNV) cells, mean ± SD (n=3). Statistical analysis shown above the plot, n.s., not significant.

### G3BP1 colocalizes with viral replication complex during MNV infection

Viral replication complexes (RC) of RNA viruses are observed as cytoplasmic foci characterized by the co-localisation of non-structural proteins required for the replication of the genomic RNA and viral dsRNA replication intermediates. Single cell analysis of the subcellular distribution of dsRNA in MNV-infected BV2 GFP-G3BP1 cells at different times post-infection showed a strong colocalisation of both NS3 and dsRNA in the juxtanuclear structures previously shown in Figure 2, further defining them as potential MNV replication centers (Figure 4A). Remarkably, those structures also contain GFP-G3BP1 and are discernable as early as 6h p.i. and at 9h p.i, suggesting the recruitment of G3BP1 to MNV replication complexes is concomitant with the accumulation of viral products. Of note, the redistribution of G3BP1 to this juxtanuclear structure does not occur in arsenite-treated cells indicating that this accumulation is a viral-induced event rather than a ubiquitous response to stress in this cellular model as suggested by Figure 2A.

**Figure 4.**
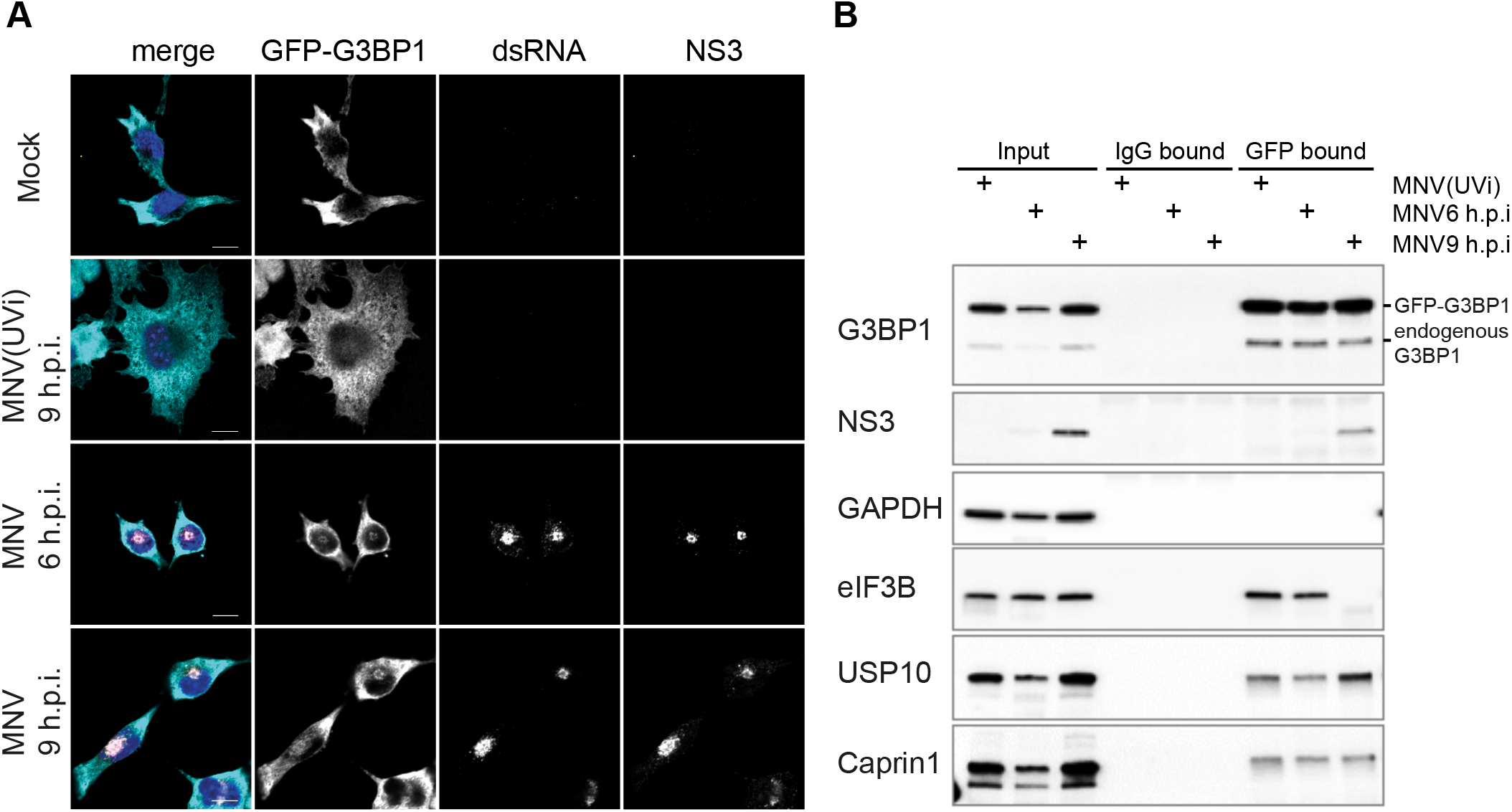
G3BP1 colocalises with MNV replication complex. GFP-G3BP1 colocalises *in vivo* with dsRNA and NS3 in MNV-infected cells (A). Mock-, MNV(UV)-or MNV-infected BV2 GFP-G3BP1 cells (MOI 10) were incubated for 6 and 9h p.i prior fixation. Representative view (n=2) of a confocal analysis of the subcellular distribution of GFP-G3BP1 (cyan), showing MNV replication complexes identified by double immunodetection against dsRNA (magenta) and MNV NS3 (gold). Nuclei were stained with DAPI. Scale bars, 10μm. GFP-G3BP1 co-precipitates *in vitro* with MNV NS3 (B). Representative western blot analysis (n=3) of a co-immunoprecipitation assay against GFP-G3BP1 (GFP) performed in extracts of cells infected as described above. A parallel co-precipitation using a normal IgG has been used as a control (IgG). Levels of cellular and viral proteins are shown in Inputs, IgG-bound fractions and GFP-bound fractions.

To further investigate a potential interaction between MNV products and G3BP1, we performed an immunoprecipitation assay from MNV-infected cells lysates. As G3BP1 is a protein containing several different motifs of interaction with proteins and nucleic acids (22,63) and known for its ability to homo- and hetero-polymerize, we addressed G3BP1 interacting partners in BV2 cells expressing GFP-G3BP1 and used an anti-GFP antibody rather than a G3BP1 antibody, allowing us to capture entire G3BP1-containing complexes in cells infected with MNV or MNV(UVi), at 6 or 9h p.i. As shown in Figure 4B, the immunoprecipitation of GFP-G3BP1 led to a strong enrichment of G3BP1 in the GFP-bound fractions for all three conditions but not for the control IgG-bound fractions. The absence of GAPDH in the anti-GFP bound fractions confirms the specificity of the experimental conditions. Paralleling the co-localisation of NS3 and G3BP1 seen in fixed cells by confocal microscopy (Figure 2), we observed substantial co-immunoprecipitation of NS3 in the anti-GFP bound fraction at 9h p.i. with MNV. Interestingly, analysis of the known interactors of G3BP1 revealed different patterns of coprecipitation with no significant differences between the MNV(UVi) and the MNV-infected cells for Caprin1 and USP10, and a striking absence of interaction at 9hpi for the translation initiation factor eIF3B (Figure 4B) and other translation factors (data not shown). Noteworthy, in the absence of viral stress, G3BP1 is able to interact with its known stress granule partners USP10, Caprin1 or eIF3B, as previously reported by interactome studies (16,66), supporting the existence of networks of RBP interactions in the cytoplasm that help nucleate granules when later required. Overall, this supports the previously observed absence of recruitment of eIF3B with G3BP1 aggregates in MNV-infected cells, and their resistance to CHX treatment observed by confocal microscopy (Figure 2), which could suggest the assembly virus-induced granules is independent from the exchange of mRNPs with polysomes.

### MNV-induced G3BP1 granules differ from canonical SG

SGs are dynamic membrane-less structures that undergo fusion, fission and move in the cytosol (67). In addition to mRNA, RNA-binding proteins and translation factors, different subsets of SGs have been shown to contain specific proteins or mRNAs associated with the specific pathways resulting in their assembly, *i.e* components of the interferon (IFN) signaling pathway during infection or subsets of RBPs during neurodegeneration-associated stress (13). Recently, an increased understanding of SG biogenesis and function has been achieved by characterizing the RNA and protein composition of SGs in yeast and mammalian cells (16,17,66). These studies relied on affinity enrichment or proximity ligation using G3BP1 as a bait to unravel the SG interactome. Therefore, we reasoned these could be adapted to characterise MNV-induced G3BP1 granules. To this end, BV2 GFP-G3BP1 cells were either treated with 0.1 mM arsenite for 1h to induce SG assembly or infected with MNV for 9h. G3BP1 granules and SG cores were then enriched by sequential centrifugation and purified by immunoprecipitation using antibodies to GFP (to trap GFP-G3BP1) or IgG (as a control) followed by pull down with Protein A-conjugated Dynabeads as previously described (68). Epifluorescence microscopy analysis then confirmed the isolation on anti-GFP beads of SG core or G3BP1 granules, while no granules could be detected in the control IgG immunoprecipitation (Figure 5A). To characterize the identity of G3BP1-interacting partners within these, mass spectrometric analysis was performed.

**Figure 5.**
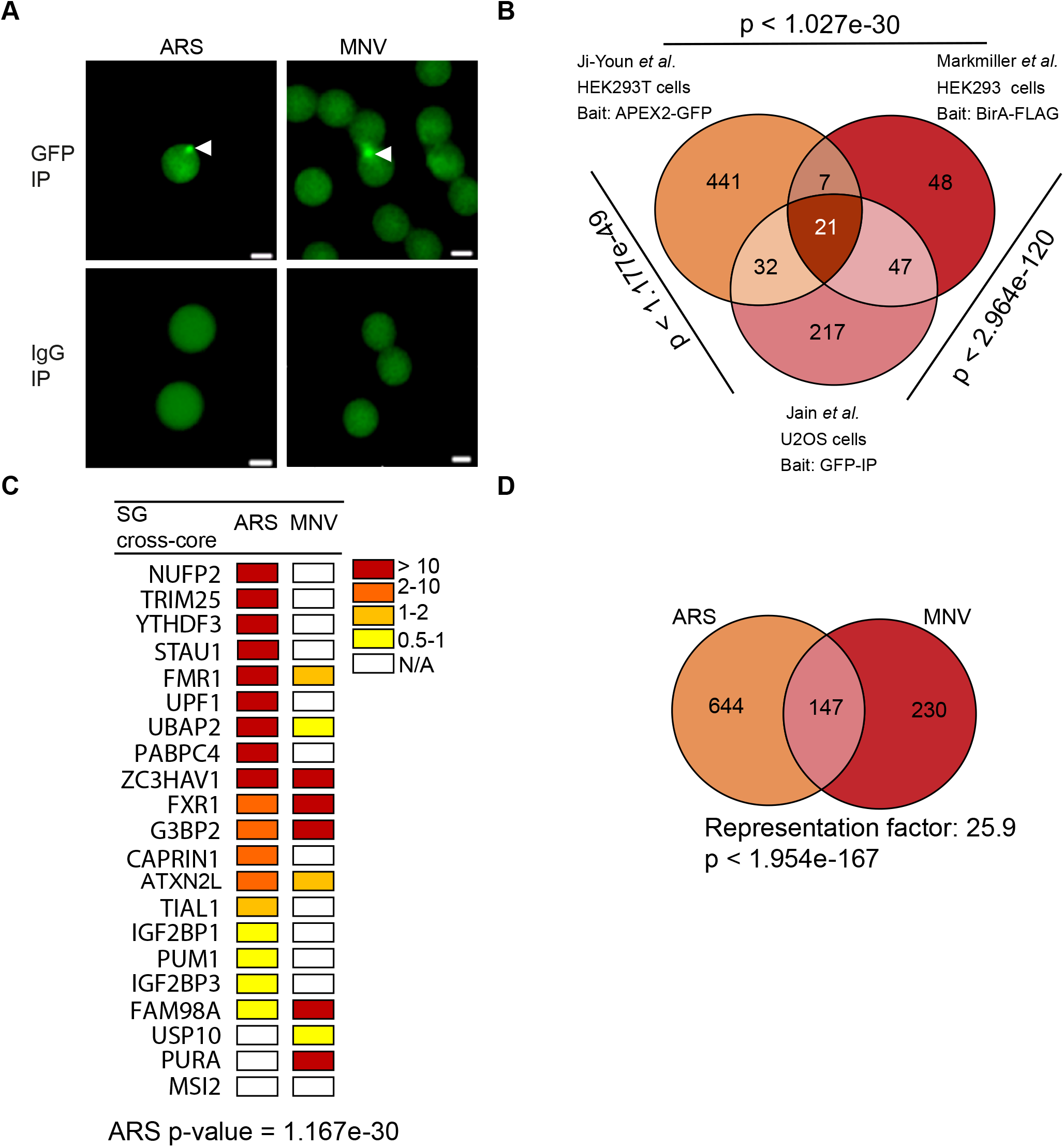
MNV-induced G3BP1 granules are distinct from arsenite-induced stress granules. (A) Dynabeads bound to GFP and to IgG analysed by epifluoresence microscopy in arsenite-stressed and in MNV-infected cells; bead-bound G3BP1 granules are indicated by white arrows (B) Venn diagram of the SGs interactome showing the common elements between three different stress granules (SG) interactome identified in human cells. The hypergeometric p-value is displayed by the side of each group. (C) Heat map representing the 21 proteins identified as “SG cross-core” in arsenite-treated and MNV-infected G3BP1 interactome ranked by the Log2 fold changes. Fold changes are indicated with from red to yellow colour bar and white for N/A. (D) Venn diagram between 795 proteins identified in ARS and 379 in MNV infected cells. The representation factor showed the enrichment over the expected and p-value is based on the cumulative distribution function (CDF) of the hypergeometric distribution of the data set over the mouse proteome.

Mass spectrometry analysis identified 791 proteins from arsenite-treated cells and 387 proteins from MNV-infected cells using for the filtering criteria ≥1 Log2-fold changes of MS peak areas of immunoprecipitated proteins compared to the respective IgG antibody (Table S1). First, to ensure that we had successfully isolated SGs, we compared the proteins identified in arsenite-treated BV2 cells (mouse) to those identified in human cells using a similar procedure (17). While this revealed that 66 out of the 791 (this study) or 317 (Jain *et al)* proteins significantly overlap between their compositions (Figure S3; Table S2; overlap between the two list p < 3.234e-55, hypergeometric test), analysis of the enriched proteins reveals that arsenite-induced SG in murine cells are significantly enriched, >10 log2 folds above background, in previously identified components of the SG core such as translation factors (*i.e* eIF2Balpha, eIF2Bepsilon, eIF4E and PABP2), RNA binding proteins (*i.e* PUM3, FMR1, NUFP2, FUS), tRNA synthetases (*i.e* tyrosyl-tRNA synthetase 2, YARS2), ribosomes biogenesis (*i.e* BRX1 and Rrs1), ATPases (*i.e* ATP2B, ATP1A3, RAD50) and RNA or DNA helicases (*i.e* Ddx47, Helz2 and ATRX) (Figure S4, Table S3). Next, we generated a refined list of common SG proteins identified with different experimental approaches, GFP-based immunoprecipitation (17), BirA-FLAG or APEX2-GFP for proximity labelling (16,66) but all using G3BP1 as bait. We defined this list as “SG cross-core proteins” (Table S4). The overlap analysis identified 21 proteins which are consistently and significantly enriched by the different experimental strategies across the three studies (Figure 5B). From this list of SG cross-core proteins, we were able to identify a significant enrichment of 18 out of 21 proteins in our arsenite-induced SG proteins (Figure 5C, p value = 1.167e-30). With confidence that we had successfully isolated SG in arsenite-treated BV2 cells, we next compared the G3BP1 granulome generated in MNV-infected cells. Strikingly, although a significant subset of proteins was enriched in both the treatment (147 proteins, p value= 6.999e-174), we identified 230 proteins enriched in MNV infected samples and not in the ARS-treated cells; and 644 proteins in arsenite-only (Figure 5D and table S2). Comparison of the mammalian SG proteome and G3BP1 granulome in infected cells clearly identified that different families of proteins are enriched (Table S3). For example, ribosomal RNA processing like RRP1, RRP8, RRP9, Ddx17, Ddx47, Ddx51 and Ddx27, are found in arsenite-treated cells while BST2 which is an IFN-induced antiviral host restriction factor (69) or guanyl-nucleotide exchange factors like Dock11 and Dock2 in response to chemokine signaling (70,71) are found in MNV-infected cells and not arsenite-treated cells (Table S3). These results altogether clearly support a model in which MNV infection result in a redistribution of G3BP1 cellular protein partners within granules, which may contribute to counteracting the assembly of SGs during infection.

### MNV-induced translational stalling is independent from the cellular stress response

The absence of canonical SG accumulation during MNV infection, despite a marked translational shut-off and increased eIF2α phosphorylation led us to hypothesize an uncoupling between the P-eIF2α-dependent stress pathways and translation suppression. The inhibitor of the Integrated Stress Response Inhibitor (ISRIB) has previously been shown to reverse the inhibitory impact of P-eIF2α on its recycling factor eIF2B, allowing the exchange between GDP and GTP on phosphorylated substrates, thereby rescuing translation and resolving stress (72,73). To address the downstream activation of P-eIF2α pathway in MNV-infected cells, we measured the effect of ISRIB on translation shut-off, MNV replication and G3BP1 localisation. As expected, treatment of the cells with 200nM ISRIB resolved the accumulation of stress granules induced by arsenite in BV2 GFP-G3BP1 cells (Figure 6A and C). Stimulation of RAW264.7 cells with tunicamycin, a known ER-stress inducer which activates the kinase PERK (74), resulted in a translation shut-off, also reverted in ISRIB-treated cells (Figure 6B and D). In marked contrast, ISRIB treatment was unable to rescue the translational shut-off observed in MNV-infected cells at 10h p.i. (Figure 6B and D) and had no impact on the accumulation of G3BP1 foci during MNV infection (Figure 6A and C). Finally, treatment with increased concentrations of ISRIB from 1 to 1000nM had no impact on viral replication as measured by TCID50 assays (Figure S5). These results suggest an uncoupling between eIF2α phosphorylation and translation suppression during MNV infection, and supports the assembly of virus-specific G3BP1 foci rather than SGs.

**Figure 6.**
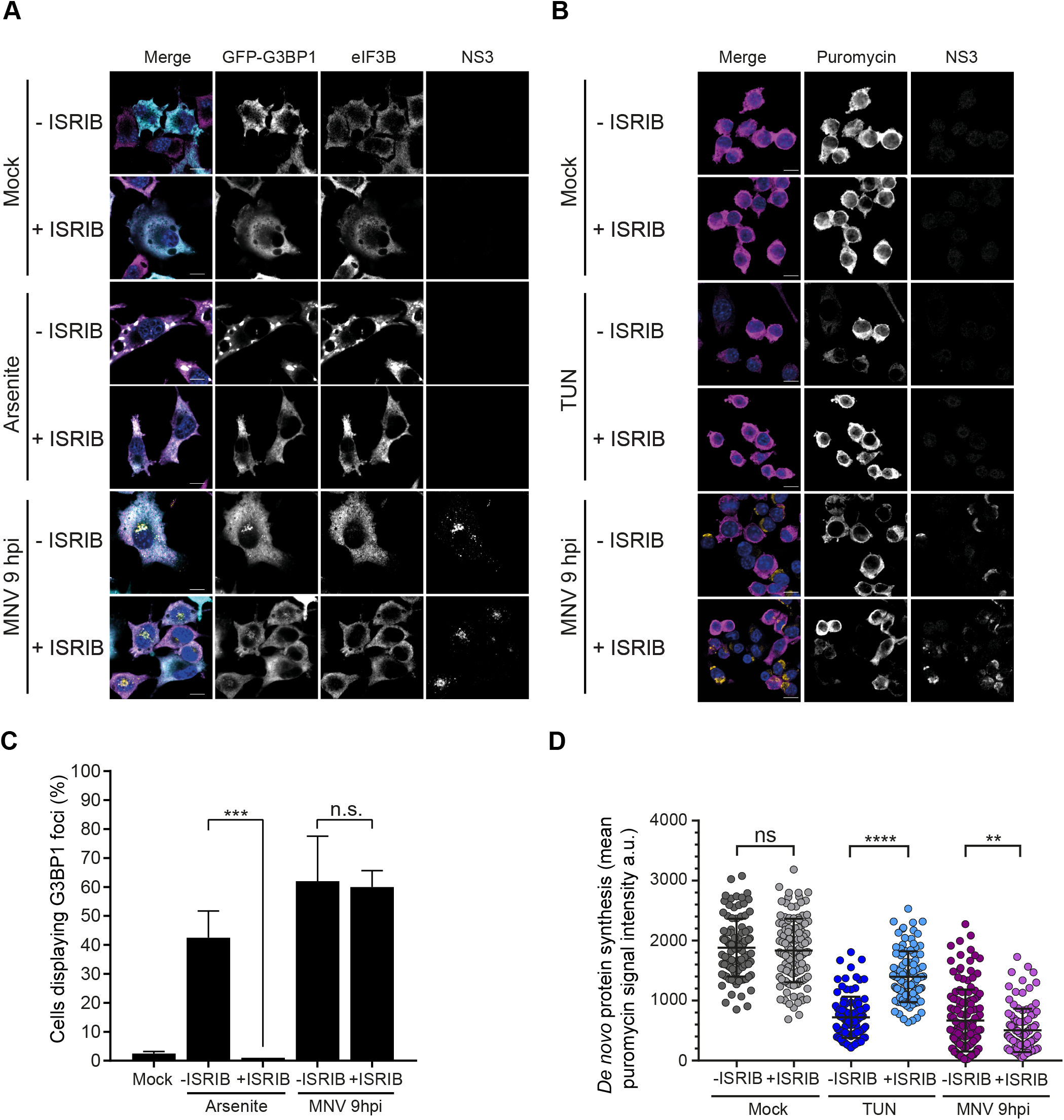
The P-eIF2α signaling inhibitor ISRIB has no effect on G3BP1 granules assembly nor on translation shut-off in MNV-infected cells. MNV-induced G3BP1 granules are insensitive to ISRIB treatment (A and C). BV2 GFP-G3BP1 cells were infected with MNV (MOI 10) for 9hp.i. or treated with 0.1mM of arsenite (ARS) for 45 min, with (+ISRIB) or without (-ISRIB) 200nM of ISRIB. Arsenite-treated cells were used as positive control for SG formation. (A) Representative views of confocal analysis (n=3) of the subcellular distribution of GFP-G3BP1 (cyan), counterstaining of stress granules by immunodetection of eIF3B (magenta) and detection of the infected cells by immunostaining against MNV NS3 (gold). Nuclei were stained with DAPI. Scale bars, 10μm. (C) Bar plots of the number of cells displaying GFP-G3BP1 foci as a percentage of the GFP-positive cells (Mock and arsenite) or GFP- and NS3-positive cells (MNV), mean ±SD (n=3). Statistical significance given above the bars, ***, *P* <0.001, n.s., no significance. ISRIB treatment does not rescue MNV-induced translation shut-off (B and D). RAW264.7 cells were infected with MNV (MOI 10) for 10hp.i. or treated with 5μg/ml of tunicamycin (TUN) for 6h, with (+ISRIB) or without (-ISRIB) 200nM of ISRIB. Tunicamycin-treated cells were used as positive control for P-eIF2α-dependent translation shut-off. Cell cultures were incubated with 10μg/ml of puromycin to label the nascent polypeptidic chains. Puromycin-labelled chains were visualised by immunostaining against puromycin (magenta) and infected cells were detected by immunostaining against MNV NS3 (gold). (B) Representative views of the confocal analysis. Scale bars, 10μm. (D) Representative scatter plots of *de novo* protein synthesis measured by fluorescence intensity of the puromycin signal (n=3), a.u. arbitrary units. Statistical significance and number of analysed cells (n) are given at the top. **** *P* < 0.0001, **, *P* < 0.01, n.s., no significance.

### MNV infection leads to a metabolic-induced phosphorylation of eIF2α

To further understand the importance of eIF2α-dependent pathways during MNV infection, we engineered MEF cells expressing wild-type (wt) or a non-phosphorylatable mutant of eIF2α (MEF S51A, (75)) to make them susceptible to MNV infection by constitutively expressing the MNV receptor CD300lf (40). The accumulation of viral proteins by immunoblotting at 10h p.i. confirmed the ability of MNV to replicate in these cells (Figure 7A). Furthermore, the ability of MNV to produce infectious particles showed no difference between the wt and S51A MEF as measured by viral yield assay (Figure 7B), suggesting that MNV replication is independent from eIF2α phosphorylation. Given that both the regulation of eIF4E and PABP activities were previously shown to contribute to MNV-induced translational shut-off (44,53), we postulated this it could occur independently from eIF2α phosphorylation. To address this, protein synthesis was quantified using ribopuromycylation assays in MEF expressing S51A. When compared to mock, infection with MNV at 10h p.i. resulted in the expected reduction in global protein synthesis activity (Figure 7C and D). This confirmed that the phosphorylation of eIF2α during MNV infection is uncoupled from the observed translational shut-off.

**Figure 7.**
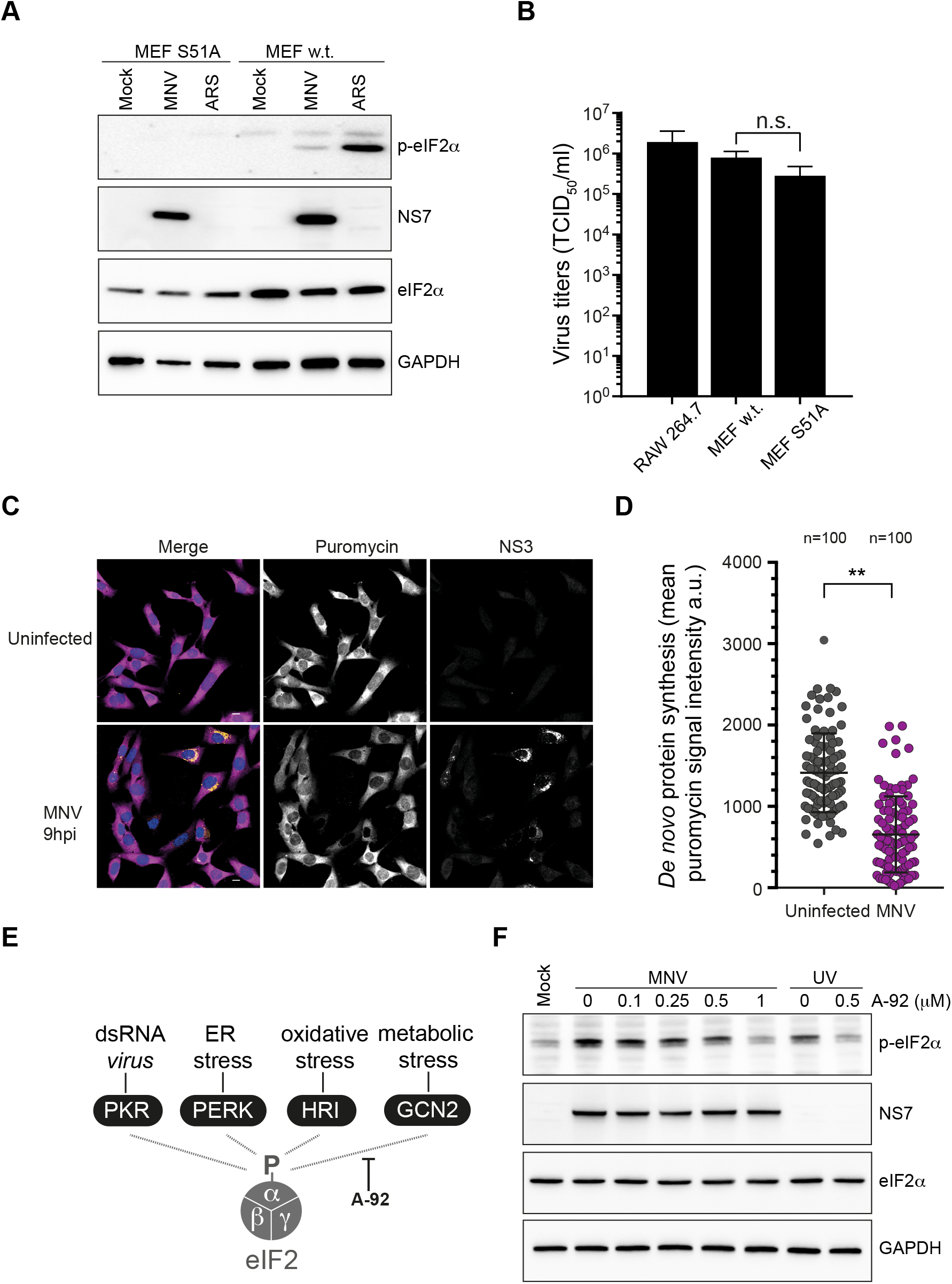
MNV-induced P-eIF2α is uncoupled from translational shut-off and is dependent upon GCN2 activity. P-eIF2α is dispensable for MNV infection (A and B). Wild type (w.t.) or expressing unphosphorylatable eIF2α (S51A) MEF cells, constitutively expressing MNV receptor CD300lf were infected with MNV (MOI 10) for 10hp.i. Arsenite-treated cells were used as control. (A) Representative western blot analysis (n=2) of the translation of MNV NS7 in absence of P-eIF2α. P-eIF2α has no effect on MNV replication (B). Bar plots of the viral titre measured by TCID50 (logarithmic scale) from MEF(w.t.) and MEF(S51A) infected with MNV (MOI 1) for 16h. RAW264.7 cells were used as control. Mean ±SD (n=3), statistical analysis given above the bars, n.s, not significant. MNV-induced translation shut-off is independent of P-eIF2α (C and D). Uninfected (Mock) or MNV-infected MEF(S51A) cells (MOI 10) for 10h p.i. were incubated with 1μg/ml of puromycin to label the nascent polypeptidic chains. Puromycin-labelled chains were visualised by immunostaining against puromycin (magenta) and infected cells were detected by immunostaining against MNV NS3 (gold). (C) Representative views of the confocal analysis, scale bars, 10μm. (D) Representative scatter plots (n=2) of *de novo* protein synthesis measured by fluorescence intensity of the puromycin signal of uninfected (-MNV) and MNV-infected (+MNV) cells, a.u. arbitrary units. Statistical significance and number of analysed cells (nj are given at the top. ** *P* < 0.01. MNV-induced P-eIF2α is dependent upon GCN2 activity (E and F). (E) Diagram of the pathways leading to the phosphorylation of eIF2α and showing the inhibition of GCN2 by A-92. (F) Representative western blot analysis (n=2) of RAW264.7 cells, naïve (Mock) or infected with MNV for 10hp.i. (MOI 10) in presence of increasing doses of A-92 ranging from 0.1μM to 1μM (MNV). UV-irradiated RAW264.7 cells were used as control, cultivated in parallel and treated with 0.5μM of A-92 (UV).

Next, we interrogated the dynamic quantitative nature of eIF2α phosphorylation during MNV infection. To this end, we used Phos-tag acrylamide gel electrophoresis on a time course infection of RAW264.7 cells by MNV or MNV(UVi), allowing a direct quantification of the phosphorylated form compared to the total level of the protein using the same antibody on the same gel (49,50). As shown in Figure S6, eIF2α is identified mainly as in two forms, the lower molecular mass product corresponding to the non-phosphorylated form and the weaker upper one corresponding to P-[S51]-eIF2α. While no significant changes in eIF2α phosphorylation were observed during infection with MNV(UVi), infection with MNV resulted in a gradual increase in the amount of P-eIF2α from 3.53% to 11.37% at 10h p.i., compared to a maximum of 17.94% in arsenite-treated cells. Interestingly, this reveals that only a modest fraction of the total eIF2α pool needs to be phosphorylated to result in the strong accumulation of SGs associated with arsenite-treatment, and that MNV infection results in comparable, albeit slightly lower, eIF2α phosphorylation level than the strong oxidative conditions linked to arsenite stimulation.

During infection the recognition of the viral dsRNA replication intermediate of RNA viruses by the recognition pattern receptors and eIF2α kinase PKR is expected to trigger non-self sensing and eIF2α phosphorylation (9). Yet, the absence of correlation between eIF2α phosphorylation and translational efficiency, led us to question which of the four known kinases, all activated by different stresses, drives eIF2α phosphorylation during MNV infection (Figure 7E). In response to metabolic stresses such as UV irradiation and nutrients depletion, the kinase GCN2 has been shown to induce a phosphorylation of eIF2α as well as a P-eIF2α-independent translation shut-off (76,77). We therefore investigated the role of GCN2 activity on MNV-induced phosphorylation of eIF2α using the GCN2 inhibitor A-92 (78). Increasing concentrations of A-92 added at 0h p.i. to MNV-infected RAW264.7 cells resulted in a dose-dependent inhibition of eIF2α phosphorylation, reduced to background levels (Figure 7F) confirming the origin of this phosphorylation. Overall these results suggest that the MNV-induced translational shut-off occurs independently from eIF2α phosphorylation, which itself is driven by GCN2 activation rather than non-self PKR sensing.

## DISCUSSION

In the present study, we have addressed how murine norovirus, a positive strand RNA virus, evades the host anti-viral response in macrophages cell lines and immortalized cells by investigating step-by-step and challenging the activation of the ubiquitous axis P-eIF2α – SG assembly. Despite what could be seen as manifest hallmarks of induction of this classical anti-viral response during MNV infection (Figure 1), we have demonstrated the absence of canonical SG formation and, oppositely, the assembly of MNV-specific granules enriched in G3BP1 (Figure 2). Moreover, proteomic analysis of MNV-induced granules showed a composition drastically different from arsenite-induced SG in BV2 cells (Figure 5), further demonstrating the unique nature of such aggregates.

In addition to the known levels of control by post-translational modifications, the ability of G3BP1 to nucleate SGs has been linked to its interaction with either Caprin1 or USP10, where USP10 plays an inhibitory anti-aggregation role (63). This interaction involves FGDF motifs on Caprin1 and USP10, competing for the NTF2 motif on G3BP1. The switch between USP10 and Caprin1 binding modifies the physical properties of G3BP1, allowing interaction with translationally inactive mRNA through binding to the 40S subunits of stalled initiating transcripts and culminates by the nucleation of SGs by LLPT (63). Notably, a mimicry of the inhibitory G3BP1-USP10 interaction has been described during infection by old world alphaviruses, preventing the formation of SGs by G3BP1 sequestration and involving FGDF motifs in the viral protein nsp3 (79). No such motifs had been identified in MNV proteins and as human and murine G3BP1 NTF2 motifs are identical, this seems to rule out a similar mechanism of G3BP1 sequestration during MNV infection. Furthermore, we observed that only a fraction of the cellular G3BP1 seems aggregated and redistributed during MNV replication, suggesting that enough G3BP1 would be available for SG assembly (Figure 2 and 4A). Also, we demonstrated that G3BP1 extracted from MNV-infected cells is physically able to undergo a LLPT and to form SG-like aggregates (Figure 3). Thus, the condensation of G3BP1 in those MNV-induced granules does not seem responsible for preventing SG formation by sequestering G3BP1, as opposed to what happens during alphavirus replication.

Rather intriguingly, a study aiming at the discovery of the host factors required for MNV replication in murine macrophages cell line BV2 by CRISPR screening showed the requirement of G3BP1 for MNV to achieve a productive replication, suggesting a viral repurpose of this host anti-viral factor (40). In the present work, we have observed an unexpected spatio-temporal colocalisation of G3BP1 granules with MNV replication complexes (RC) by confocal microscopy (Figure 2 and 4A) and a molecular interaction between G3BP1 and viral factors by immunoprecipitation (Figure 4B). The observed pattern of aggregation of G3BP1 fits the subcellular localization of MNV RC described as associated with membranes derived from the secretory pathway and proximal to the microtubule organizing center (56–58). Interestingly, while MNV infection does not abrogate the ability of G3BP1 to coalesce given the right physico-chemical conditions, no viral factors have been found associated with the *in vitro* condensed SG-like aggregates (Figure 3 and data not shown) suggesting that the association of G3BP1 with the MNV RC does not result from molecular crowding and LLPT. Finally, the time-dependent increasing recruitment of a fraction of the available cellular G3BP1 concomitant with the accumulation of viral products in the RC (Figure 4A) thus evokes the hijacking of this factor by a stoichiometric type of interaction with either viral proteins or nucleic acids and supports the concept that G3BP1 recruitment to MNV RC could be a potential requirement to the viral replication.

To further investigate whether eIF2α phosphorylation had a role in G3BP1 granules formation and recruitment into MNV RC, we used the inhibitor of the integrated stress response ISRIB which had been shown to reverse the downstream effects of phosphorylation, namely accumulation of stalled initiating transcripts and SG accumulation. By this chemical inhibition of P-eIF2α downstream signaling (Figure 6 A and C) and using a cellular model expressing a non-phosphorylatable form of eIF2α (Figure 7 A-D), we showed an uncoupling between the phosphorylation of eIF2α, the formation of MNV RC and G3BP1 recruitment to this structure. Importantly, this observation also correlates with an LLPT-independent formation of G3BP1 granules and an LLPT-independent interaction between G3BP1 and viral factors in MNV-infected cells, further demonstrating the absence of SG characteristics in this virus-induced G3BP1 aggregates.

Recently, the understanding of proteins involved or associated with RNA granules and stress granules in particular has exploded mainly through their biochemical purification, proximity mapping of RNA granules associated proteins or fluorescence-activated particle sorting (16,17,66,80). To further investigate the evasion of the stress granule response during MNV infection we characterized G3BP1 interactions within virus-induced granules and compared them to well-characterized arsenite-induced SGs (Figure 5). First, our analysis revealed that arsenite-induced SGs in murine cells differ in their composition to those assembled in human cells. However, we recapitulated the enrichment in proteins previously identified as SGs components using similar procedures such as translation factors, RNA binding proteins, tRNA synthetases, ribosomes biogenesis factors, ATPases, RNA or DNA helicases ((17) and Figure S4). Furthermore, combining the different SGs interactome studies that used G3BP1 as bait to generate a cross-core set of stress granules components revealed that 18 out of 21 of these proteins are enriched in arsenite-induced stress granules isolated from murine cells, confirming the conservation of the interactions across species. In contrast, during MNV infection, only 3 out of 21 of the cross-core stress granules proteins associated with G3BP1, namely ZC3HAV1, FXR1 and G3BP2. Further analysis of the G3BP1 granulome during infection clearly confirmed a shift in composition with 230 out of 377 proteins associated with G3BP1 specific to MNV-induced granules (Figure 5 and table S2). Of the 147 proteins common with arsenite-induced stress granules, enrichment revealed proteins associated with RNA transport (GO:0051028) and localisation (GO:0006403) and ribonucleoprotein complex biogenesis (GO:0022613) while proteins involved in immune response-regulating cell surface receptor signaling pathway (GO:0002433) and involved in endocytosis (GO:0006897) are associated with MNV-induced granules (table S3). Overall, while further biochemical and structural characterization of virus-induced G3BP1 granules is required to fully understand their function, our results support a model in which MNV infection results in a redistribution of G3BP1 cytoplasmic partners, which could provide the basis for evasion of stress granules assembly during infection.

In contrast to the dogmatic understanding of P-eIF2α signaling leading to accumulation of stalled initiating transcripts and assembly of SG, we observed an uncoupling between P-eIF2α and the inhibition of translation in MNV-infected cells (Figure 6B and D, Figure 7C and D). This could explain the absence of SG formation as only stalled initiation transcripts have been shown crucial for SG assembly (81). Moreover, MNV infection and arsenite treatment led to similar levels of eIF2α phosphorylation suggesting that there should be enough P-eIF2α to inhibit eIF2B and regulate the host initiation of translation (Figure S5), which further supports a viral antagonism strategy at the eIF2α signaling nexus.

Noticeably, it has been reported that some forms of stress lead to a phosphorylation of eIF2α correlating with only moderate translation shut-off and without any SG formation (76,77). The authors identified the eIF2α kinase GCN2, potentially activated downstream of the translation shut-off itself and remarkably reversing the causality relationship between P-eIF2α and translation inhibition. By analogy, we challenged the eIF2α response in MNV-infected cells using the novel inhibitor of GCN2 A-92 (78) and demonstrated that the phosphorylation of eIF2α is indeed dependent upon GCN2 activity (Figure 7F). Because viruses are obligate intracellular parasites, it is thus tempting to speculate that the translation shut-off occurs as a result of a cellular status of starvation likely generated by the exponential and unchallenged synthesis of viral proteins, rather than through sensing of intermediates of replication and genomic RNA. Furthermore, there are few reported cases of GCN2 activation by viruses, which overall seems to restrict the viral fitness in *in vivo* studies (82). As MNV virion production does not seem to be affected by ISRIB (Figure S5) and GCN2 inhibition had no effect on the synthesis of viral non-structural protein (Figure 7F), it seems quite unlikely that the phosphorylation of eIF2α by GCN2 had any antiviral effects, at least *in vitro*, nor it being repurposed for MNV replication.

While it has been published that a non-replicative MNV RNA transfected into cells does not trigger a PKR-dependent signaling (83), our results would suggest that the amplification of dsRNA also seems to stay surprisingly unnoticed by this particular pattern recognition receptor, at least in our model. Previous electron microscopy imaging of cryosections of MNV-infected RAW264.7 cells highlighted that MNV dsRNA and the RNA polymerase NS7 are located on the cytoplasmic side of MNV-induced vesicles rather than in their lumen, which suggest a cytoplasmic replication available for recognition (56). In light of our results, several hypotheses as to the nature of MNV strategies of PKR avoidance can thus be proposed by analogy with other classical viral mechanisms, such as “hiding in plain sight” or direct inhibition of the non-self recognition machinery. As PKR has also been described as a transducer of MDA5 dependent anti-viral response beside its eIF2α kinase activity (84), this could also explain the poor interferon response to MNV infection (85). In particular, this hypothesis would fit the model where the MNV factor VF1 is involved in the antagonism of the IFN response by interaction with the MAVS axis (86) and its activation by MDA5 (87,88). Remarkably, it has also been shown that human norovirus RNA replication does not trigger any noticeable IFN response in mammalian cells (89). It is thus fairly reasonable to suspect a common mechanism of avoidance of the host response among the *Norovirus* genera, manifested in this study by the evasion from eIF2α signaling stress responses. Furthermore our observation of a redistribution of interactions for the RBP G3BP1 paves the way for further studies investigating how MNV impacts on the global landscape of RBP interactions during infection and how these contribute to viral replication mechanisms.

## AVAILABILITY

Fiji is an open source image processing package based on ImageJ (http://fiji.sc/wiki/index.php/Fiji). Metascape is an open source web portal for gene annotation and analysis resources (http://metascape.org/gp/index.html#/main/step1). Bioinformatics statistical analysis were computed using CGI scripts hosted by Nematode bioinformatics (http://nemates.org/MA/progs/representation.stats.html). Cytoscape is an open source software platform for visualizing complex networks (https://cytoscape.org/).

## ACCESSION NUMBERS

The mass spectrometry proteomics data have been deposited to the ProteomeXchange Consortium via the PRIDE (90) partner repository with the dataset identifier PXD011956.

## Supporting information

## SUPPLEMENTARY DATA

Supplementary Data are available at NAR online.

## ACKNOWLEDGEMENT

The authors wish to thank the members of the Locker laboratory for providing feedback on the manuscript.

