## Supplementary material for "Norovirus infection results in assembly of virus-specific G3BP1 granules and evasion of eIF2α signaling"

**SUPPLEMENTARY MATERIAL AND METHODS**

**Mass Spectrometric analysis**

Samples were run into an SDS-PAGE gel for approximately 1cm and the gel lane was then excised and subjected to in-gel tryptic digestion using a DigestPro automated digestion unit (Intavis Ltd.). The resulting peptides were fractionated using an Ultimate 3000 nano-LC system in line with an Orbitrap Fusion Tribrid mass spectrometer (Thermo Scientific). In brief, peptides in 1% (vol/vol) formic acid were injected onto an Acclaim PepMap C18 nano-trap column (Thermo Scientific). After washing with 0.5% (vol/vol) acetonitrile 0.1% (vol/vol) formic acid peptides were resolved on a 250 mm x 75 μm Acclaim PepMap C18 reverse phase analytical column (Thermo Scientific) over a 150 min organic gradient, using 7 gradient segments (1-6% solvent B over 1min, 6-15% B over 58min, 15-32%B over 58min, 32-40%B over 5min, 40-90%B over 1min, held at 90%B for 6min and then reduced to 1%B over 1min) with a flow rate of 300 nL min−1. Solvent A was 0.1% formic acid and Solvent B was aqueous 80% acetonitrile in 0.1% formic acid. Peptides were ionized by nano-electrospray ionization at 2.2 kV using a stainless steel emitter with an internal diameter of 30 μm (Thermo Scientific) and a capillary temperature of 250°C. All spectra were acquired using an Orbitrap Fusion Tribrid mass spectrometer controlled by Xcalibur 2.0 software (Thermo Scientific) and operated in data-dependent acquisition mode. FTMS1 spectra were collected at a resolution of 120 000 over a scan range (m/z) of 350-1550, with an automatic gain control (AGC) target of 400 000 and a max injection time of 100ms. The Data Dependent mode was set to Cycle Time with 3s between master scans. Precursors were filtered according to charge state (to include charge states 2-7), with monoisotopic precursor selection and using an intensity range from 5E3 to 1E20. Previously interrogated precursors were excluded using a dynamic window (40s +/-10ppm). The MS2 precursors were isolated with a quadrupole mass filter set to a width of 1.6m/z. ITMS2 spectra were collected with an AGC target of 5000, max injection time of 50ms and HCD collision energy of 35%. The raw proteomic mass spectrometry data files were processed using Proteome Discoverer software v1.4 (Thermo Scientific) and searched against the Uniprot Mouse database using the SEQUEST algorithm. Peptide precursor mass tolerance was set at 10ppm, and MS/MS tolerance was set at 0.6Da. Search criteria included carbamidomethylation of cysteine (+57.0214) as a fixed modification and oxidation of methionine (+15.9949) as a variable modification. Searches were performed with full tryptic digestion and a maximum of 2 missed cleavage sites were allowed. The reverse database search option was enabled and all peptide data was filtered to satisfy false discovery rate (FDR) of 1%. Following this, proteins represented by at least two peptides, in at least two of the three technical replicates with PE=1 (PE=ProteinExistence), were selected, and the peptides from the antibody heavy chain were removed from the analysis. The criteria applied for the normalisation were based on peak area where each protein area is represented as a fraction of the total protein area for the whole sample and before in order to avoid zero values an arbitrary value (0.000001) was added to all peak areas. The calculation of the fold changes between each paired IP-GFP over the corresponding IP-IgG was performed and then the average of the three technical replicates was transformed into a log2. The Gene Ontology annotation (GO) was performed using Metascape (http://metascape.org/gp/index.html#/main/step1) which identified all statistically enriched terms referring to GO, KEGG, REACTOME, CORUM databases and calculates the accumulative hypergeometric p-values and enrichment factors. The significant overlap between different groups was calculated using hypergeometric probability formula: (http://nemates.org/MA/progs/representation.stats.html) "C(D, x) * C(N-D, n-x) / C(N,n).

**SUPPLEMENTARY TABLES, FIGURES AND MOVIES LEGENDS**

Supplementary Figure S1. MNV replication is not affected by exogenous expression of SG markers in BV2 cells. (A) Bar plots of the viral titres measured by TCID50 (logarithmic scale) from BV2 cells w.t., Puro, Neo, Puro-Neo, GFP-G3BP1, mCherry-eIF3E and GFP-G3BP1/mCherry-eIF3E inoculated with MNV (MOI 1) for 16h. Mean ±SD (n=3), statistical analysis given above the bars, n.s, not significant. BV2 GFP-G3BP1 cells were infected with MNV for 9hp.i prior fixation. (B) Representative view of confocal analysis (n=2) of GFP-G3BP1 subcellular localisation with immunodetection of G3BP1 (magenta) and MNV NS3 (gold). Scale bars, 10µm.

Supplementary Figure S2. MNV-induced G3BP1 aggregation is observed in living cells. Representative view of a time-lapse acquisition by confocal microscopy of BV2 cells expressing GFP-G3BP1 (cyan) and mCherry-eIF3E (magenta) in culture infected with MNV (MOI 20) at 10h15 p.i. Scale bar, 5µm.

Supplementary Figure S3. Analysis of stress granules components between mouse and human cells. Venn diagram of the SGs interactome showing the common elements between human cells (U2OS cells) and mouse cells (BV2 cells). The hypergeometric p-value and enrichment factor are displayed.

Supplementary Figure S4. GO analysis of stress granules components in mouse cells. Cytoscape clustering was performed using ClueGO app based on GO terms (molecular function and biological process) considering two side hypergeometric test and Bonferroni correction p-value<0.005, using GO term fusion and layout "perfused force direct" for a clear representation. The nodes were grouped accordingly to GO terms and the node size corresponds to the significance of each GO term in the network. We arbitrarily added an additional colour pattern to the cluster in order to highlight the difference between the two groups on analysis (ARS and MNV).

Supplementary Figure S5. The P-eIF2$\alpha$ signaling inhibitor ISRIB has no effect on MNV replication. Bar plots of the viral titres measured by TCID50 (logarithmic scale) from BV2 cells inoculated with MNV (MOI 1) for 16h in presence of increasing doses of ISRIB from 10nM to 1µM. Mean ±SD (n=3), statistical analysis given above the bars, n.s, not significant.

Supplementary Figure S6. Quantitative analysis of P-eIF2$\alpha$ abundance in MNV-infected RAW264.7 cells by PhosTag. Representative PhosTag acrylamide gel analysis (n=3) of RAW264.7 cells infected (MOI 10) for the indicated times with replicative MNV (upper panel) or non-replicative virus MNV(UV) (lower panel). Naive cells (Mock) were cultivated in parallel for 10h and arsenite-treated cells were used as control. Lower band, unphosphorylated P-eIF2$\alpha$, upper band, P-eIF2$\alpha$. The percentages of

P-eIF2α compared to total eIF2α for each sample are given below the blots, Total eIF2α a being the sum of the signal intensity of the lower and upper bands in each lane.

Supplementary table S1. List of G3BP1 interactome identified by Mass spectrometry analysis in BV2 cells. List of the proteins for the 3 biological replicates from arsenite-treated cells (A) and from MNV-infected cells (B). The filter criteria applied is ≥1 Log2-fold changes of average MS peak areas of respective proteins of the immunoprecipitation over the respective IgG antibody.

Supplementary table S2. List of proteins used to plot the Venn diagram in Supplementary Figures S3 and for Supplementary Figure 4.

Supplementary table S3: Meta-analysis of pathways enrichment of G3BP1 granulome in arsenite-induced and MNV infected stress granules. Table of the list of the GO, KEGG, REACTOME and CORUM terms with significant hypergeometric p-value with associated gene name belonging to each category. (A) GO analysis performed from the whole granulome identified in BV2 cells upon arsenite treatment. (B) GO analysis performed from 230 proteins specifically identified in BV2 cells upon MNV infection not present in the granulome upon arsenite stress.

Supplementary table S4. List of G3BP1 interactome identified from three independent studies: (i) from Ji-Youn et al. the list of 501 proteins was obtained applying an arbitrary cut-off of ≥ 1 Log2 fold change between the (counts in the purification divided by counts in the controls); from Jain et al the list of 317 proteins was selected considering ≥2 fold change spectral counts of stressed over not stressed cells; (iii) from Markmiller et al 123 proteins were identified in HEK293 cells from Log2 H/L ratio.

Supplementary Movie S1: MNV-induced G3BP1 aggregation is observed in living cells. Representative view of a time-lapse acquisition by confocal microscopy of BV2 cells expressing GFP-G3BP1 (cyan) and mCherry-eIF3E (magenta) in culture infected with MNV (MOI 20), 0.5 fps, from 9h30 to 11hp.i., scale bars 10µm.

**Supplementary Figure 1**

A

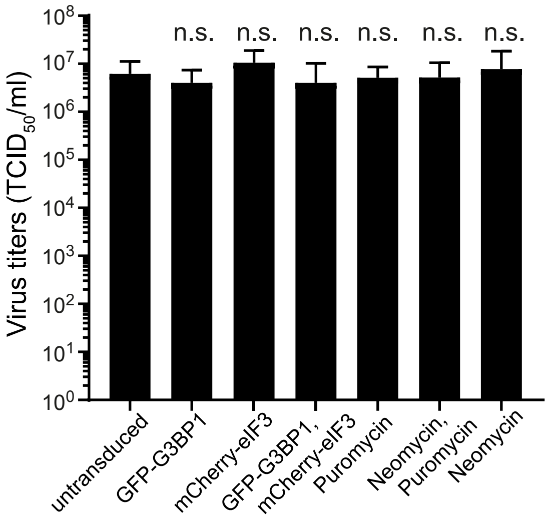


B

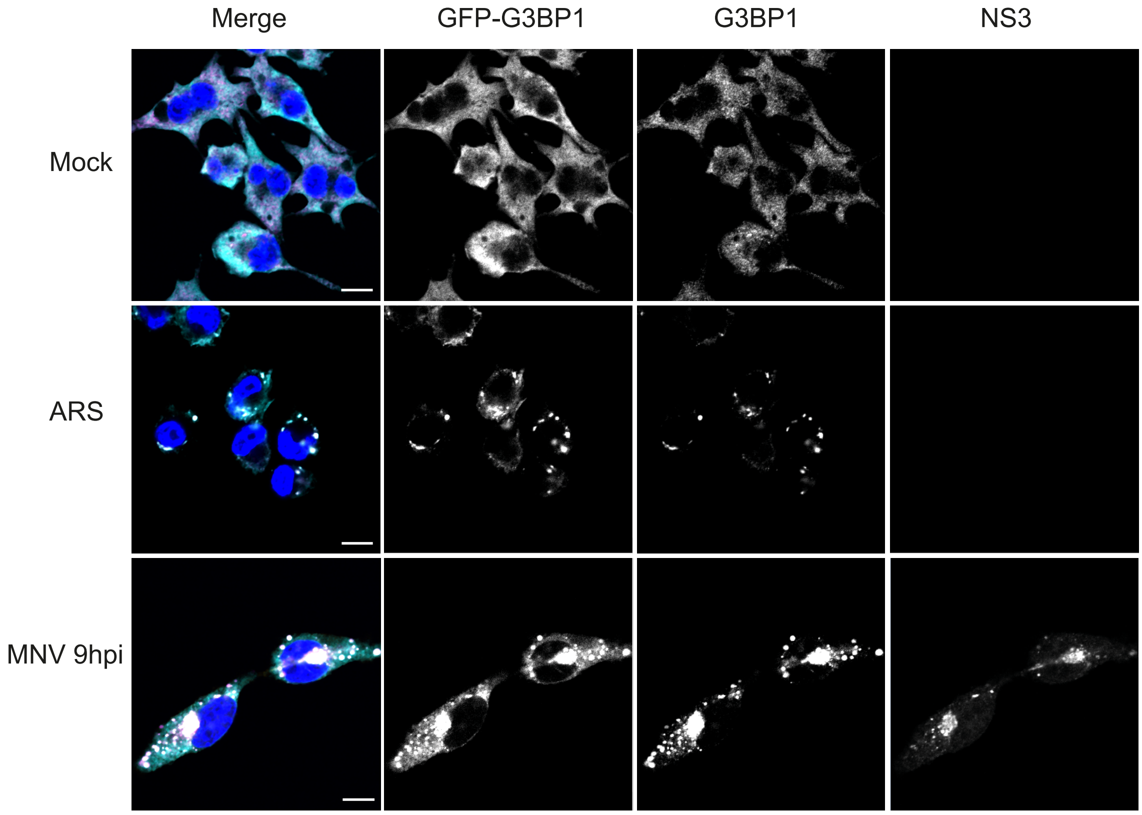

**Supplementary Figure 2**

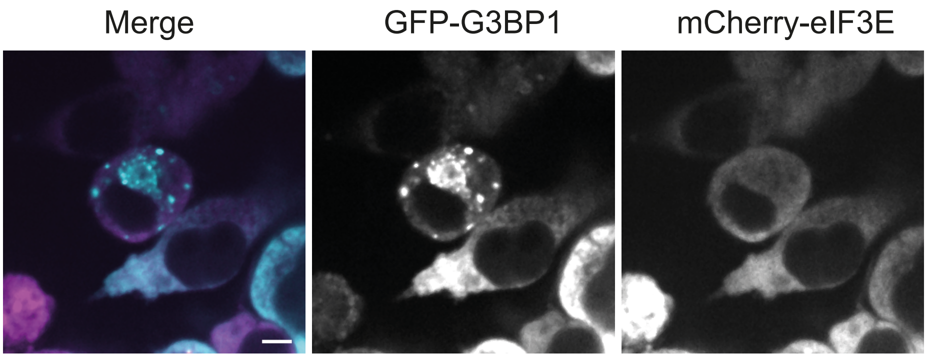

**Supplementary Figure 3**

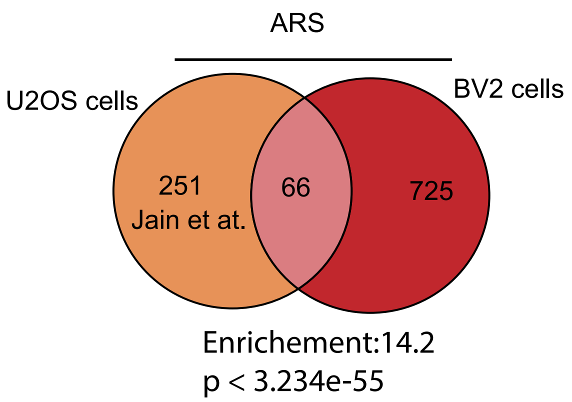

### Supplementary Figure 4

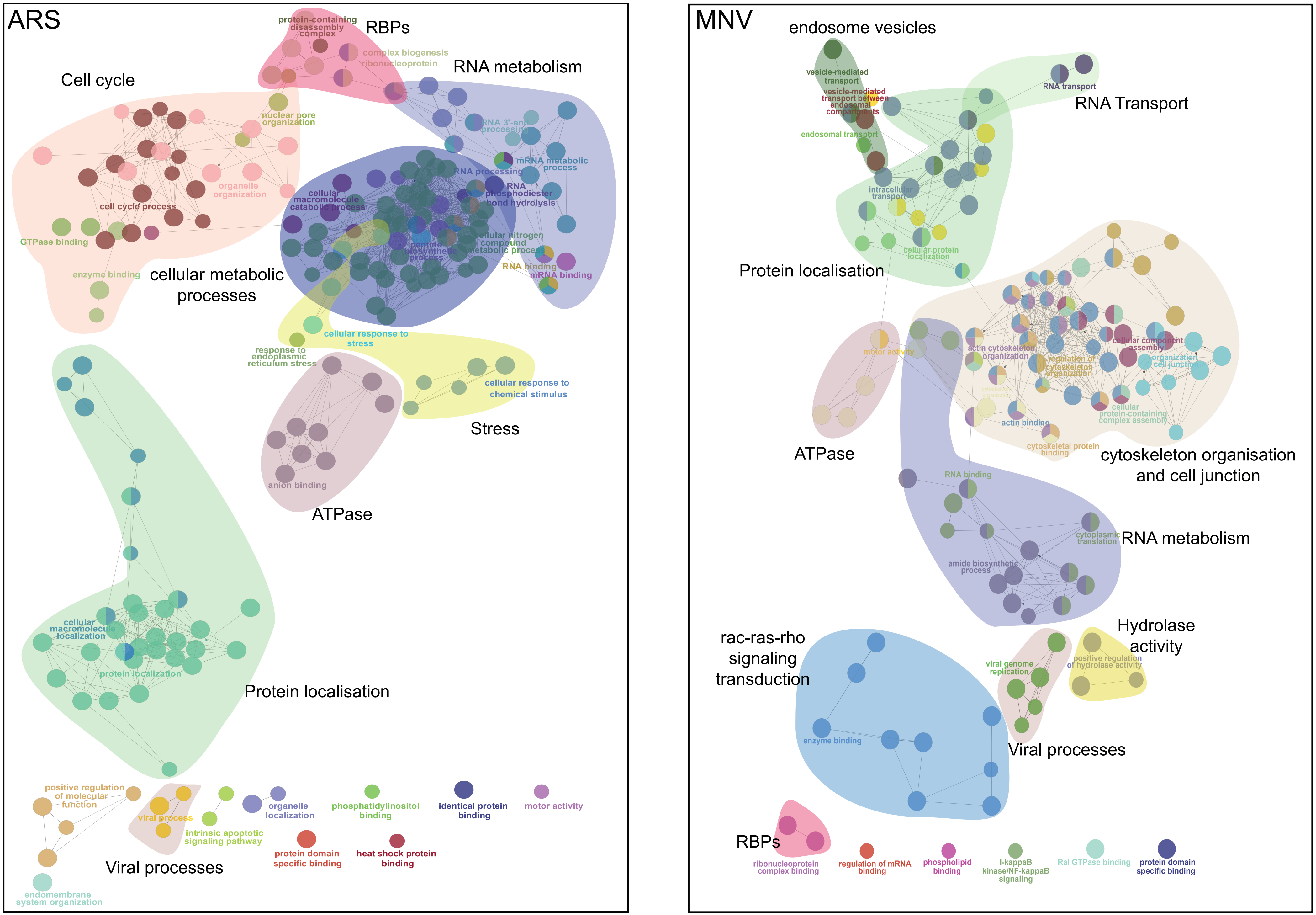

### Supplementary Figure 5

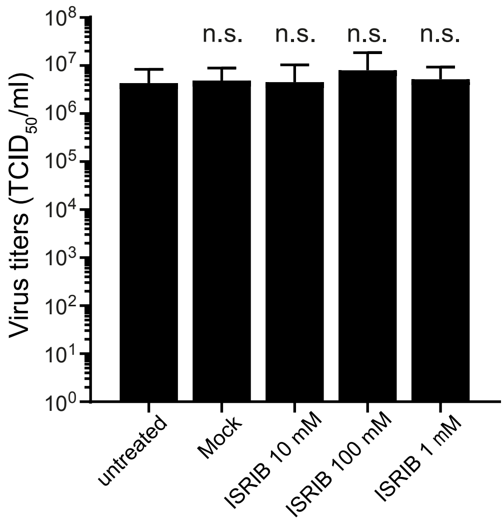

**Supplementary Figure 6**

**A**

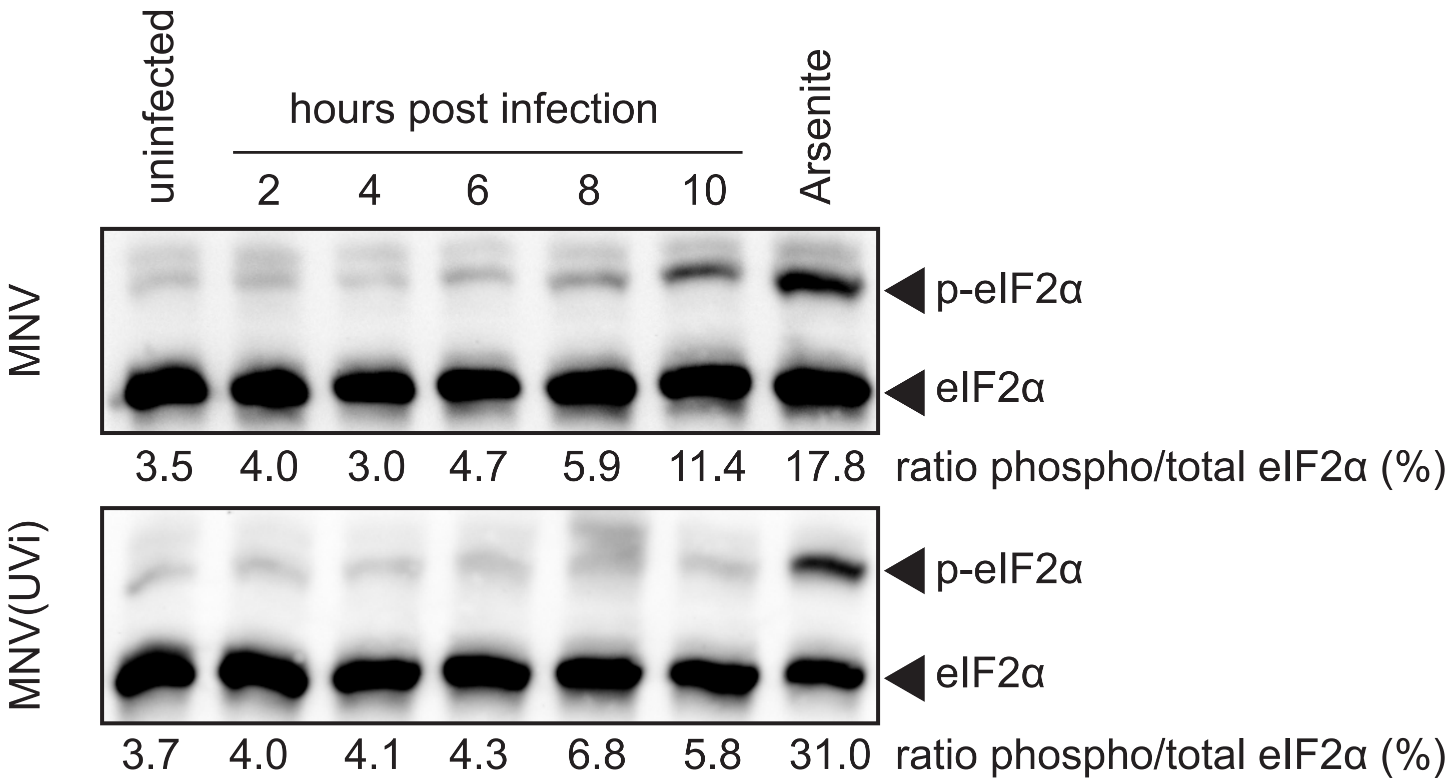


**B**

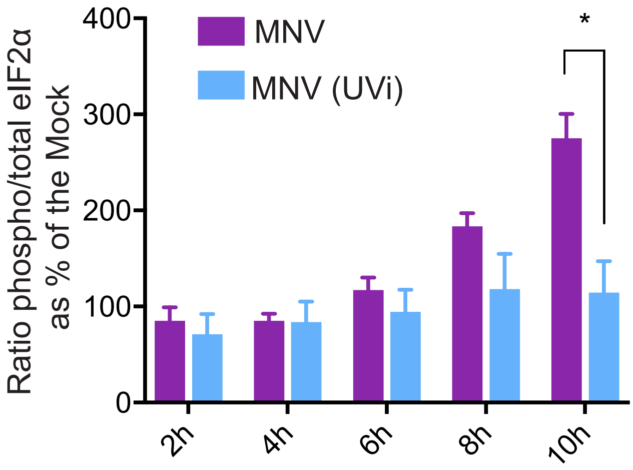
